# Mapping the Cellular Biogeography of Human Bone Marrow Niches Using Single-Cell Transcriptomics and Proteomic Imaging

**DOI:** 10.1101/2024.03.14.585083

**Authors:** Shovik Bandyopadhyay, Michael Duffy, Kyung Jin Ahn, Minxing Pang, David Smith, Gwendolyn Duncan, Jonathan Sussman, Iris Zhang, Jeffrey Huang, Yulieh Lin, Barbara Xiong, Tamjid Imtiaz, Chia-Hui Chen, Anusha Thadi, Changya Chen, Jason Xu, Melissa Reichart, Vinodh Pillai, Oraine Snaith, Derek Oldridge, Siddharth Bhattacharyya, Ivan Maillard, Martin Carroll, Charles Nelson, Ling Qin, Kai Tan

## Abstract

The bone marrow is the organ responsible for blood production. Diverse non-hematopoietic cells contribute essentially to hematopoiesis. However, these cells and their spatial organization remain largely uncharacterized as they have been technically challenging to study in humans. Here, we used fresh femoral head samples and performed single-cell RNA sequencing (scRNA-Seq) to profile 29,325 enriched non-hematopoietic bone marrow cells and discover nine transcriptionally distinct subtypes. We next employed CO-detection by inDEXing (CODEX) multiplexed imaging of 18 individuals, including both healthy and acute myeloid leukemia (AML) samples, to spatially profile over one million single cells with a novel 53-antibody panel. We discovered a relatively hyperoxygenated arterio-endosteal niche for early myelopoiesis, and an adipocytic, but not endosteal or perivascular, niche for early hematopoietic stem and progenitor cells. We used our atlas to predict cell type labels in new bone marrow images and used these predictions to uncover mesenchymal stromal cell (MSC) expansion and leukemic blast/MSC-enriched spatial neighborhoods in AML patient samples. Our work represents the first comprehensive, spatially-resolved multiomic atlas of human bone marrow and will serve as a reference for future investigation of cellular interactions that drive hematopoiesis.

## Introduction

The bone marrow is a complex organ which houses diverse cells from the hematopoietic, mesenchymal, endothelial, vascular smooth muscle and neural lineages^1^. Relatively rare non-hematopoietic cells are known to make critical contributions to physiologic functions including bone formation, immune regulation, endocrine function, and support of hematopoiesis - in particular through hematopoietic stem and progenitor cells (HSPCs) interacting with the bone marrow niche^2^. The concept of a hematopoietic niche and its constituents, first proposed 45 years ago by Schofield^3^, has been a rapidly evolving and often controversial topic^4,5^. Many cell types, including endothelial cells, perivascular stromal cells, and osteoblasts have been proposed to be critical constituents of the hematopoietic stem cell (HSC) niche^6–9^, and functional approaches have been deployed to investigate which of these elements are critical for hematopoiesis. Most of these studies relied on genetic reporters using Cre-recombinase tied to a cell-type specific promoter^10^, or are focused on one specific microenvironmental cell type. However, recent technological advances employing single-cell RNA sequencing (scRNA-Seq)^4,11–13^ revealed the existence of multiple subpopulations of non-hematopoietic marrow elements in mice, enabling better understanding of the native, unbiased cellular composition of the bone marrow niche.

Despite extensive work studying single human hematopoietic cells (e.g., Human Cell Atlas - Immune Cell Atlas), there remains a relative paucity of analogous studies defining the non-hematopoietic cells that make up the human bone marrow microenvironment^14–16^. Strictly defining the single-cell composition of the bone marrow niche has been limited by challenges in isolating sufficient viable non-hematopoietic cells, which are thought to make up <0.5% of the marrow cellularity^17,18^ and thus not captured without enrichment. One important study used cell sorting to overcome this limitation in bone marrow aspirate samples where the non-hematopoietic frequency was 0.002%, finding that mesenchymal stromal cells (MSCs) were largely homogenous in marrow from healthy donors^19^. However, specific isolation methods may greatly influence the diversity of cell types captured, as first shown in mice^12^. Bone marrow aspiration, analogous to flushing of mouse bones, may fail to capture cells which are tightly adhered to the bone surface and bias the MSC composition towards adipocytic, rather than osteogenic cells. Additionally, much of human MSC profiling has been performed following a period of *ex vivo* expansion^20,21^, which may alter the transcriptional profile of MSCs from their *in vivo* homeostatic state. Furthermore, most aforementioned functional and transcriptomic analysis of bone marrow niche cells lacked information on the spatial context of these cells.

Here, we overcome these limitations by performing multiomic profiling of single cells from healthy bone marrow samples from fresh femoral head tissue without the need for expansion. We employed scRNA-Seq to comprehensively characterize and define the transcriptomic profiles of both hematopoietic and non-hematopoietic cell types directly from human bone marrow, and to delineate patterns of intercellular communication and the mediating signaling molecules. In parallel, we used Co-Indexing by Epitopes (CODEX)^22^ to image human bone marrow with single-cell spatial proteomic resolution and performed unsupervised analysis of bone marrow niches and their cell type composition. For the first time, we uncover the full non-hematopoietic diversity in the human bone marrow niche and quantitatively define how niche elements organize hematopoiesis. Furthermore, we demonstrated the utility of our new atlas in acute myeloid leukemia (AML) patients, where we discovered enriched MSC-leukemia spatial interactions. Collectively, this work represents the first comprehensive, spatially-resolved multiomic human bone marrow atlas which will serve as a reference for future studies of the human bone marrow niche.

## Results

### A Comprehensive scRNA-Seq Atlas of the Human Bone Marrow

To uncover the cellular diversity of the human bone marrow ecosystem, we developed an experimental pipeline for enzymatic release of cells from femoral head tissue(Supplemental Figure S1A-B). We confirmed the areas of interest had normal trabecular structure by performing micro-computed tomography (micro-CT) analysis of the bone volume/trabecular volume ratio which was 0.261 +/-0.098 (n=6, Supplemental Figure S1A). Next, we devised a stepwise magnetic enrichment strategy that would capture hematopoietic cells (RBC depletion) as well as rare stem and progenitor cells (CD34 enrichment) and non-hematopoietic microenvironmental cells (CD45 depletion) (Figure 1A, see Materials and Methods). The three populations were pooled in a ratio favoring representation of rare HSPCs and non-hematopoietic cells and subjected to scRNA-Seq.

**Figure 1.**
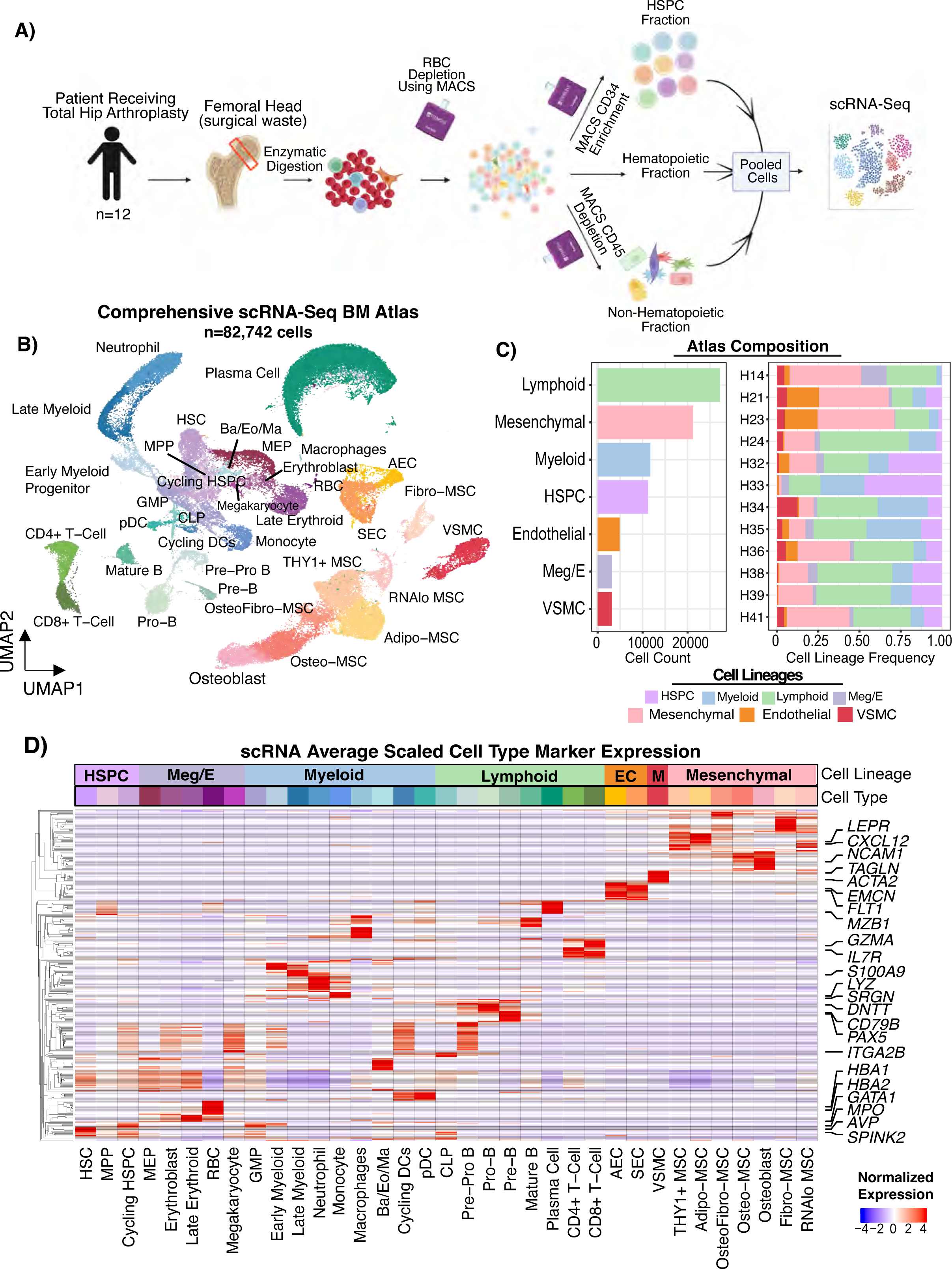
A Single-Cell transcriptomic atlas of hematopoietic and non-hematopoietic cells of human bone marrow. **A**) Schematic describing the sample collection and cell isolation strategy. Magnetic-activated cell sorting (MACS) separation of hematopoietic, stem/progenitor, and mesenchymal fractions were performed and then pooled into one scRNA-Seq reaction per patient. **B**) UMAP representation of 82,742 single-cell transcriptomes from bone marrow of 12 individuals. AEC, arterial endothelial cell. SEC, sinusoidal endothelial cell. VSMC, vascular smooth muscle cell. Ba, Basophil. Eo, Eosinophil. Ma, Mast Cell. RBC, red blood cell. pDC, plasmacytoid dendritic cell. CLP, common lymphoid progenitor. MEP, megakaryocyte erythroid progenitor. GMP, granulocyte monocyte progenitor. MPP, multipotent progenitor. HSPC, hematopoietic stem and progenitor cell. HSC, hematopoietic stem cell. Meg/E, megakaryocyte/erythroid. MSC, mesenchymal stromal cell. **C**) Bar plots showing the cell counts for each lineage captured (left) and the per sample cell lineage frequencies normalized by the total cells in each sample (right). **D**) Heatmap with normalized gene expression scaled by column (cell type) showing the gene expression of key cell type marker genes. Rows were clustered such that genes expressed in similar cell types cluster together. The top two most significant genes by adjusted p-value comparing each cell type to all other cell types were plotted. Selected marker genes for each lineage were labeled. EC, endothelial cell. M, vascular smooth muscle.

We collected fresh femoral heads from 15 individuals who underwent hip replacements, performing scRNA-Seq on 12 of them (Supplemental Table S1). In total, we sequenced 53,417 hematopoietic cells and 29,325 non-hematopoietic cells from 12 femoral head samples (Figure 1B-C). We captured rich, high-quality transcriptional information in each cell lineage (Supplemental Figure S1C), with a per-cell dataset average of 16,903 UMIs, 3,117 unique genes, and only ∼3% mitochondrial reads detected after filtering. Importantly, we did not observe a strong inflammatory response signature, except in cell types classically associated with inflammation (mature neutrophils and monocytes), supporting the notion that these samples represent healthy marrow (Supplemental Figure S1D). Cells were subjected to unsupervised clustering and annotated based on manual examination of known marker gene expression (Figure 1D, Supplemental Table S2). As expected, we identified the hematopoietic hierarchy from HSCs (*AVP, SPINK2*) to mature hematopoietic cells of the myeloid, megakaryocyte/erythroid, and lymphoid lineages (Figure 1D, Supplemental Table S2). Additionally, we improved upon other hematopoiesis-focused bone marrow references^14,15^ by capturing the full granulocyte differentiation trajectory by virtue of our sample processing not including density-based isolation of mononuclear cells – a standard clinical tissue banking process which results in granulocyte loss due to their multi-lobated nuclei (*CSF3R*, Supplemental Figure S1E-F). Numerous plasma cells were also captured, as a byproduct of the CD45 depletion procedure (*MZB1*, Supplemental Figure S1E-F).

Next, we analyzed the non-hematopoietic cell compartment. Compared to previous single-cell references, our experimental approach revealed significant cellular diversity within the non-hematopoietic cells of the bone marrow (Supplemental Figure S1F). Within the non-hematopoietic cells, we identified three major lineages – vascular smooth muscle (*ACTA2*, *RGS5, TAGLN*), endothelial cells (*CDH5*, *PECAM1, VWF*), and mesenchymal cells (*CXCL12, PDGFRA)* including osteolineage cells and mesenchymal stromal cells (MSCs) (Figure 1B-D, Supplemental Table S2, Supplemental Figure S1E). Contrary to previous results from human bone marrow aspirate samples^19^ which found that MSCs were relatively homogenous and expressed an adipocytic transcriptional profile^23^, we observed that mesenchymal stromal cells from digested bone marrow were highly heterogenous with numerous clusters identified (Figure 1B, Supplemental Figure 1F).

### Non-Hematopoietic Cell Subset Analysis Demonstrates MSC and Endothelial Cell Transcriptional Diversity

To better understand the heterogeneity of non-hematopoietic cells, we employed unsupervised subclustering to study the endothelial and MSC clusters. Of note, prior to performing this analysis we removed a cluster of rare cells (termed RNAlo MSC) with very low unique gene and UMI counts (6.6% of mesenchymal cells) (Supplemental Figure S2A). We were unable to determine whether these cells were a true cell type or a technical artifact as they expressed low number of genes and transcripts per gene similar to low-quality cells, but were also similar in QC profile to known highly specialized cell types such as neutrophils or RBCs. We found significant heterogeneity among mesenchymal cells and identified osteo-lineage cells (*NCAM1, SPP1, BGLAP)*, adipo-lineage cells (*APOE, LPL, PPARG, CEBPA*), and fibroblastic cells (*PDPN*, *CSPG4*, *DCN*, *DPT*) (Figure 2A-B, Supplemental Figure S2B). Osteo-lineage cells were further divided into *IBSP/BGLAP*-high osteoblasts and *IBSP/BGLAP*-low Osteo-MSCs (Figure 2B). While adipocytes themselves were not captured in our dataset, likely due to losses during centrifugation steps in sample processing, we found that the cell cluster with the highest *CXCL12* level also highly expressed adipose lineage genes such as *CEBPA, PPARG*, *APOE*, and *LPL*, which we labeled as Adipo-MSC. These cells likely correspond to the previously reported murine MALPs^11^ and Adipo-CAR cells^12^. We also found a population of *THY1*+ MSCs which expressed adipocytic genes strongly, but also a unique expression profile including *THY1* and *LBP*, with lower *LPL* expression compared to Adipo-MSC which we therefore termed as THY1+ MSCs (Figure 2B, Supplemental Figure S2B). THY1+ MSCs did not have a clear murine counterpart^24^. Fibroblast-like cells which expressed *CXCL12* and very specifically expressed *PDPN*, *HAS1*, and *NT5E* were also observed, which we termed Fibro-MSC (Figure 2B). We also identified an *APOD*+ *GSN*^hi^ population of cells that had lower adipogenic gene expression (*LPL*, *LEPR*) than Adipo- or THY1+ MSCs but retained expression of *COL1A1,* clustering between Fibro-MSC and Osteo-MSC in UMAP space and thus we defined them as Osteo-Fibro MSC (Figure 2A-B). To the best of our knowledge, this population is also previously unreported in humans or mice. The Fibro-MSC population, however, had significant concordance with the Early Mesenchymal Progenitor population described previously^11^ in mice by *CPSG4*, *CD34*, and *DPT* expression, as well as to the human skeletal stem cell population^25^ defined by *PDPN*, *CD164*, and *NT5E* (CD73) expression and lack of lineage marker expression. This Fibro-MSC population also had gene expression consistent with the canonical “mesenchymal stem cell” markers defined by the International Society for Cellular Therapy^26^ (ISCT), *NT5E* (CD73), *THY1* (CD90), and *ENG* (CD105) (Figure 2C). It is important to note that expression profile of these genes was highly variable among MSC subsets, and this should be considered in future studies that attempt to isolate MSCs based on these markers. In particular, inclusion of CD90 (*THY1*) as a sorting marker may highly bias the MSC population isolated towards Fibro and THY1+ MSCs. *NES,* a marker used to label MSCs in mouse models^27^, was not detected in human MSCs but was expressed in Sinusoidal Endothelial Cell (SEC), Arterial Endothelial Cell (AEC), and Vascular Smooth Muscle Cell (VSMC) (Supplemental Figure S2B). *NGFR* (CD271) was more consistently expressed among subsets, providing rationale for its continued use^28^ as an unbiased MSC marker (Figure 2C). *MCAM* (CD146), however, was not highly expressed. It should be noted that there was considerable heterogeneity in mesenchymal cell frequency between samples, which could reflect dynamic cell states such as activation or technical factors including sampling variability.

**Figure 2.**
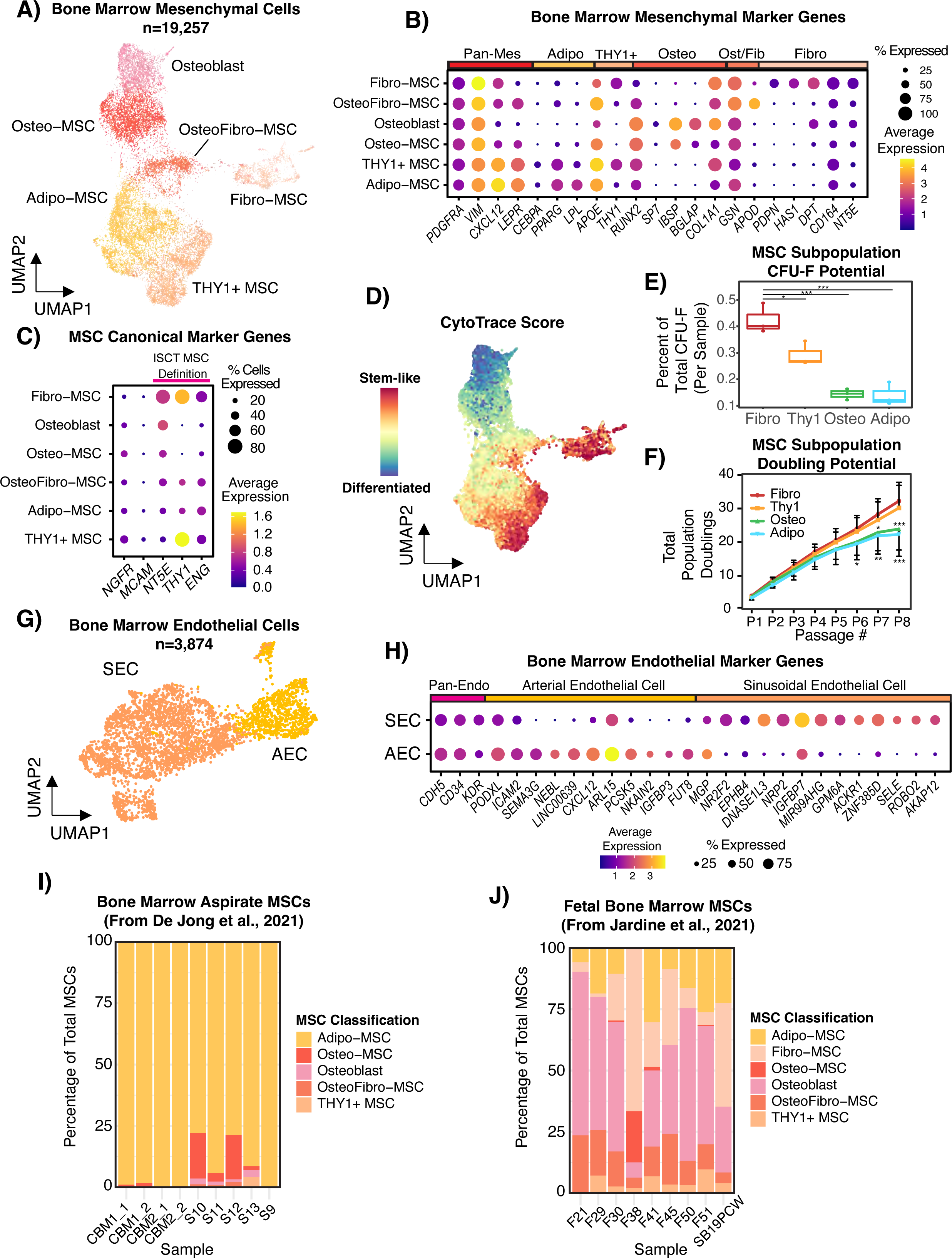
Defining the non-hematopoietic cellular composition of human bone marrow. **A**) UMAP computed from 19,257 mesenchymal cells from 12 individuals showing different mesenchymal subsets, with RNAlo MSCs being excluded. **B**) Dot plot showing normalized expression of key mesenchymal marker genes in MSC subsets. Rows and columns were manually ordered. **C**) Dot plot showing the normalized expression of literature-derived marker genes for human MSCs including *NT5E*(CD73), *THY1*(CD90), and *ENG*(CD105). *NGFR* (CD271)^28^ and *MCAM* (CD146)^72^ have also been described as canonical MSC markers. **D**) CytoTRACE analysis projected onto the MSC UMAP showing the predicted differentiation score, where higher values imply the cell is more primitive. **E)** Boxplots showing the relative colony forming potential (number of colonies produced by each cell type divided by total colonies from each sample, each data point is one sample) of sorted Fibro-MSC (CD45-CD38-CD235ab-VECAD-PDPN+), THY1+ MSC (CD45-CD38-CD235ab-VECAD-PDPN-LEPR+ CD90+), Adipo-MSC (CD45-CD38-CD235ab-VECAD-PDPN-LEPR+ CD90-), and osteolineage cells (CD45-CD38-CD235ab-VECAD-PDPN-CD56+). P-values were computed as Fibro-MSC vs. all by Dunnett’s Multiple Comparisons test. **F)** Line plot showing the population doublings over the course of cell culturing for eight passages, where all cells were passaged every 7 days. P-values are computed as Fibro-MSC vs other MSCs by two-way ANOVA. **G**) UMAP showing 3,874 endothelial cells from the 12 individuals. SEC – Sinusoidal Endothelial Cells, AEC – Arterial Endothelial Cells. **H**) Dot plot showing normalized expression of top differentially expressed genes by adjusted p-value between endothelial subsets, as well as *CDH5*, *CD34*, and *KDR* which were manually selected as pan-endothelial markers. **I**) MSCs from human bone marrow aspirate reported from De Jong et al., *Nature Immunology*, 2021 were reference mapped to our scRNA-Seq atlas and the cell type labels were predicted. **J**) Human fetal bone marrow mesenchymal cells from Jardine et al., *Nature*, 2021 were reference mapped to our scRNA-Seq atlas and the cell labels were predicted.

To explore whether any of these populations represented a more mesenchymal stem-like state, we employed unbiased pseudotime analyses using CytoTRACE^29^ to determine the developmental ordering of the cells. Fibro-MSCs were identified as the most primitive population, consistent with their transcriptomic similarity to previously described mesenchymal progenitors and the ISCT definition (Figure 2C-D). THY1+ MSCs also had a high CytoTRACE score, suggesting they may also be more primitive. To validate that Fibro and THY1+ MSCs were more stem/progenitor-like, we sorted MSC subpopulations based on our scRNA-Seq data and performed fibroblast colony forming (CFU-F) assays (Supplemental Figure S3A-3B, details in Materials and Methods). Sorted PDPN+ Fibro-MSCs exhibited the most CFU-F forming capacity, followed by THY1+ MSCs, while Osteo-MSCs and Adipo-MSCs had the lowest CFU-F forming capacity (Figure 2E). Cultured Fibro and THY1+ MSCs continued to proliferate at high passages while Adipo-MSCs and Osteo-MSCs slowed progressively (Figure 2F). Furthermore, Fibro MSCs were able to differentiate into osteoblasts, adipocytes, and chondrocytes after being cultured in the appropriate differentiation medium based on lineage specific staining and marker gene expression (Supplemental Figure 2C-D). We generally found that Adipo-MSCs were the most abundant mesenchymal cell type with a median of 32.6% (range from 3.26% to 73.3%) of the sample, while Fibro-MSC were much rarer with median 0.1% (range 0-39%), consistent with their role as an early progenitor (Supplemental Figure S2E). Taken together, our transcriptomic and functional data point towards Fibro-MSCs being consistent with the human mesenchymal stem cell defined by the ISCT and most related to previous reports of both human and murine early mesenchymal progenitors^11,25^.

To delineate the heterogeneity within bone marrow endothelial cells, we then performed subclustering analysis on *CDH5+* endothelial clusters. Our data captured two major classes of bone marrow endothelial cells – arterial endothelial cells (AECs) and sinusoidal endothelial cells (SECs) (Figure 2G). AECs were characterized by higher *CXCL12, ICAM2*, and *PODXL* expression and significantly lower expression of canonical venous genes such as *EPHB4* and *NR2F2* than SECs (Figure 2H). SECs expressed more genes classically associated with endothelial-hematopoietic cell interactions in the bloodstream, such as *ACKR1* or *SELE* (Figure 2H). We did not observe a cell type with a transcriptional signature that could be linked to Type H vessels connecting arterioles to sinusoids, which could reflect a combination of low overall cellular frequency and association of these vessels with the growth plate of young animals^30^. Additionally, we did not capture a cluster of *PROX1*+ *PDPN*+ *LYVE1*+ lymphatic endothelial cells, but did observe rare *LYVE1*-expressing cells (Supplemental Figure S2F). Similar to MSCs, there was significant variability of SEC vs. AEC frequencies per sample, although there was a clear trend towards SECs being more common (mean=67.6% with range 20-95.1%) (Supplemental Figure S2G).

These data collectively demonstrate the need to account for heterogeneous non-hematopoietic cells in studies of the human bone marrow microenvironment. To further illustrate this point, we performed unsupervised mapping of two previously published MSC datasets to our new reference, one of human bone marrow aspirate healthy MSCs^19^ and the second of otherwise healthy fetal MSCs isolated from crushed femur^31^. Our results show that bone marrow aspirate MSCs were largely Adipo-MSCs which may be less tightly anchored to the bone (Figure 2I), while fetal stroma had a strong bias for osteo-lineage cells as well as Fibro and OsteoFibro-MSCs (Figure 2J), consistent with our findings suggesting that these Fibro-MSCs represent some of the earliest mesenchymal progenitors. In sum, our atlas shows the importance of understanding the non-hematopoietic composition of different samples and will serve as a resource to better profile bone marrow non-hematopoietic heterogeneity in existing and future studies.

### MSCs, Endothelial Cells, and Osteolineage Cells Cooperate to Produce Diverse Hematopoietic Support Factors

We next asked whether heterogeneity in mesenchymal cell types translated to differential production of supportive factors for hematopoiesis. We first compared expression of selected hematopoietic niche factors which have been widely studied^2,5,32^ in non-hematopoietic cell subsets (Figure 3A). We found that different cell types specialized in production of factors in specific pathways, such as *CXCL12* being expressed the most in Adipo- and THY1+ MSCs, while *IGF1* was most expressed in Fibro-MSCs. One of the key strengths of our atlas is the simultaneous profiling of each type of niche cell, which allows comparisons of supportive factor expression intensity on a continuous, rather than binary, distribution. This enabled us to utilize CellChat^33^ to computationally interrogate predicted cellular communications in the bone marrow based on cognate ligand and receptor co-expression between two cell types (Supplemental Table S3). Predicted interactions were consistent with gene expression, as demonstrated by *CXCL12* and *SELE* interactions of MSCs and SECs with different HSPC subsets respectively (Figure 3B). Globally, mesenchymal lineage cells had the most outgoing signaling contributions (i.e. significant interactions in which they expressed the ligand), followed by endothelial cells, suggesting that these physiologically rare cell types provide important signals in the human bone marrow niche (Figure 3C).

**Figure 3.**
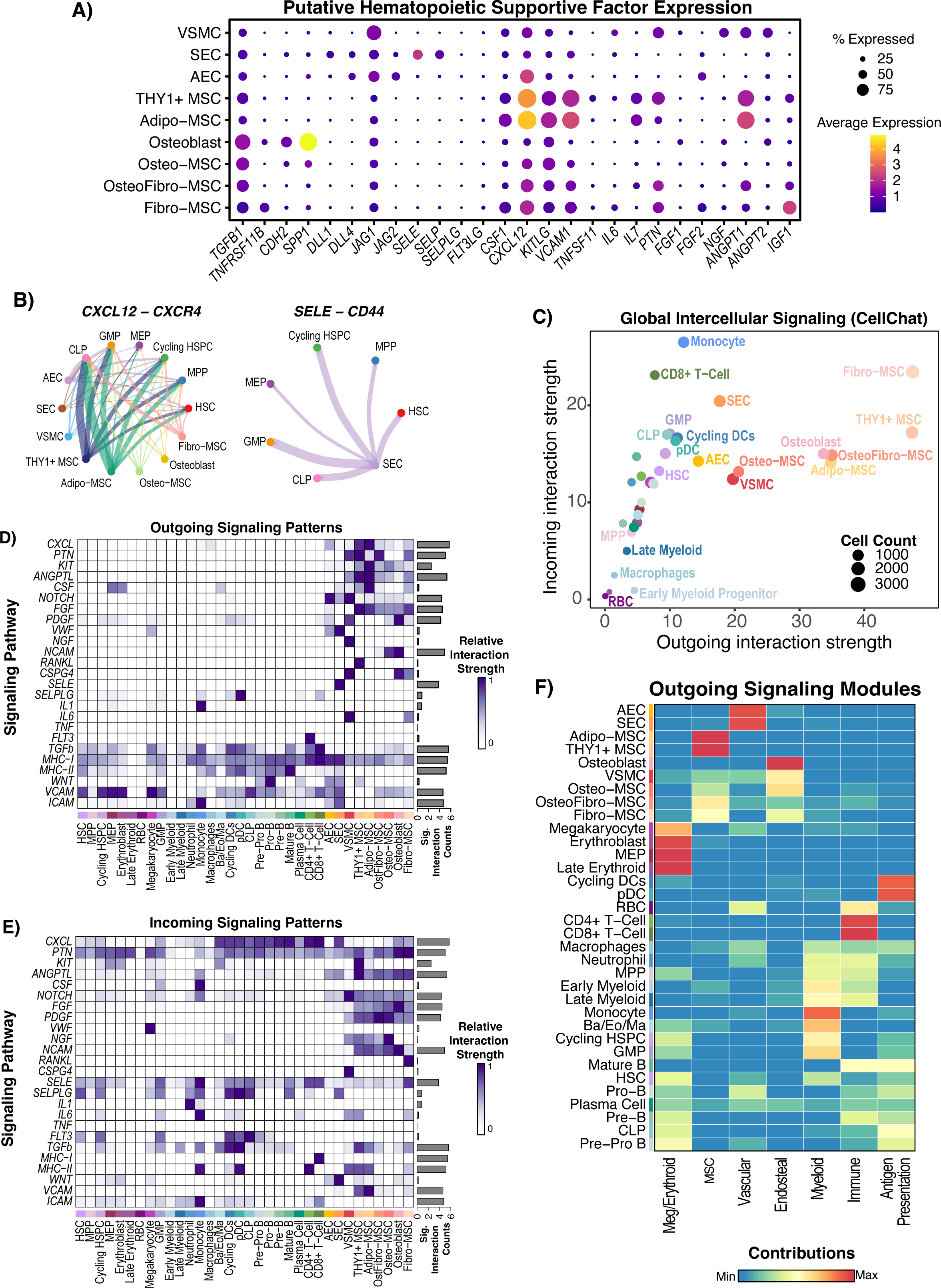
Cell-cell communication analysis reveals diverse signaling patterns between hematopoietic and non-hematopoietic cell types. **A**) Dot plot showing the expression of manually curated supportive factors for hematopoiesis in mesenchymal and endothelial populations. RNAlo MSCs were removed from the analysis as in Figure 2. **B**) Chord diagrams showing significant interactions between source mesenchymal/endothelial cell types and target HSPC and precursor cell types. Absence of a node (either source or target) means that there were no significant interactions between that cell type and the targets. Thickness of the line corresponds to the strength of the predicted interaction. **C**) CellChat network analysis with cell types scored based on their outgoing and incoming contributions to the network, where strength is defined by edge weight in the network and the count refers to the number of cells in each group. **D-E**) Outgoing (ligand enriched) and incoming (receptor enriched) significant signaling predictions were scored for each annotated signaling pathway, and the strength of signaling (edge weights) for each cell type was scaled and plotted by pathway. The barplot on the right-hand side of each panel shows the total signaling strength (network edge weights for that group of L/R pairs) across all cell groups, illustrating whether a pathway is broadly activated or not. **F**) Non-negative matrix factorization (NMF) was implemented within CellChat to identify coherent modules of outgoing signaling and the contribution of each cell type to the pattern is scored and plotted. NMF-derived signaling patterns were manually annotated based on the pathways and cell types enriched in the pattern.

Next, we interrogated the supportive factor expression and predicted communication of non-hematopoietic cells with hematopoietic cells focusing on HSPC maintenance, myelopoiesis, and lymphopoiesis. We performed predictions on a pathway level, by grouping supportive factors based on their shared pathway annotation as previously described^33^. First, we focused on HSPC maintenance. Various cell types have been reported to be important for HSPC maintenance, including osteoblasts^8,34,35^ and perivascular MSCs^6,7^. Adipo-MSCs and THY1+ MSCs produced the most well-established HSPC niche factors such as *CXCL12*^36^ (CXCL pathway) and *KITLG*^37^ (KIT pathway) which were broadly received by hematopoietic cells without strong specificity for HSPCs (Figure 3A,3D-3E). Notably, Adipo-MSC, THY1+ MSCs, and AECs expressed higher levels of, and had more predicted outgoing signaling from *CXCL12* and *KITLG* than, osteoblasts (Figure 3A, 3D). This supports mouse work which demonstrated that deletion of *CXCL12* from MSCs, but not osteoblasts, affected HSC function^6,7^ and could suggest that osteoblast CXCL12 or SCF may be less critical to support human hematopoiesis than the MSC contribution of these factors. Osteo-lineage cells did express *TGFB1 and CDH2* (Figure 3A), which have been shown to support HSC quiescence in mice^8,38^. This shows that both osteoblasts and MSCs may support human HSPCs, but most of the canonical supportive factors are produced most strongly by MSCs. Interestingly, HSCs also had significant predicted incoming signaling from E-Selectin provided by SECs (Figure 3A, 3D-3E).

We next focused on factors known to be important for myelopoiesis. *CSF1* was expressed by multiple MSC subsets, with highest expression in Adipo-MSCs. *CSF2 and CSF3*, however, were not detected at high levels, although a subset of SECs (183/3177) did have detectable expression (n.s.) of *CSF3* (Supplemental Figure S3C). This result may reflect the inherent challenge of detecting transient cytokine gene expression^39^. While most hematopoietic support factors were provided by MSCs, we also noted that CD4+ T-Cells, which are not typically thought of as critical niche cells, were the major contributors of *FLT3LG* in the bone marrow (Figure 3D). This signal was received by HSCs, GMPs, and CLPs, and is known to be critical for both myelopoiesis^40^ and lymphopoiesis^41^ (Figure 3A,3E).

We finally studied niche contributions to lymphopoiesis by interrogating factors such as *IL7* and Notch ligands, both of which are important for lymphopoiesis^42,43^. *IL7* was produced specifically by Adipo and THY1+ MSCs (Figure 3A). Broadly, AECs were characterized by an outgoing Notch signaling signature (Figure 3D). SECs expressed Notch ligands, but to a lesser degree and therefore had less overall predicted potential to engage in Notch signaling - though they expressed more *DLL1* than AECs (Figure 3A, 3D). The concept that the arteriolar niche provides the majority of Notch signaling in the bone marrow has been demonstrated in mice^4^, while in humans CD146+ MSCs^44^ have been proposed to be critical and the vascular contribution has not been clearly delineated. Collectively, endothelial cells and VSMCs are indeed the major contributors of *DLL1, DLL4*, *JAG1*, and *JAG2* in the human bone marrow, with arterioles (AEC+VSMC) expressing the most *JAG1* (Figure 3A). We also noted that Notch signaling was predicted as outgoing from mesenchymal cells (Figure 3D). Predicted mesenchymal outgoing notch signaling was weaker than endothelial notch signaling and driven largely by *JAG1* expression. Notably, all of these cell types as well as HSPCs were predicted to receive Notch signals, raising the possibility of crosstalk and coordinate regulation between niche elements.

To interrogate higher level signaling coordination, we used non-negative matrix factorization (NMF) to cluster outgoing signaling patterns based on which cells express them, resulting in modules of intercellular communication in the bone marrow. We find that different non-hematopoietic subsets contributed to different modules, which we annotated based on the cell types involved (Figure 3F, Supplemental Figure S3D). Interestingly, we found that the endosteal module was discrete from the MSC module with expression of bone-specific pathways like NGF, ncWNT, and osteopontin, showing that CXCL12-expressing MSC subsets such as Adipo-MSC and THY1+ MSC contribute globally different signals than Osteo or Fibro-MSCs, which had contributions to both the endosteal and MSC modules, while Adipo and THY1+ MSC did not contribute to the endosteal module (Figure 3F). Further, consistent with pathway analysis, AECs and VSMCs were predicted to contribute to the endosteal signaling module as well as the vascular module. Collectively, this analysis predicted interactions between bone marrow cells and their niche and shows that MSCs, vascular cells, and arterio-endosteal niches all cooperate to facilitate human hematopoiesis. Our unified atlas suggests that Adipo- and THY1+ MSCs are most important for hematopoiesis through their high expression of an MSC-specific molecular program of critical niche factors such as *CXCL12*, *KITLG*, *CSF1*, and *IL7*. Collectively, our analysis illustrates the complex intercommunicative potential of diverse human bone marrow mesenchymal and endothelial cells and that these cells are specialized to support and perhaps control different aspects of hematopoiesis.

### CODEX Multiplexed Imaging Reveals the Anatomy of the Human Bone Marrow Niche *In Situ*

We next sought to define the anatomy of both hematopoietic and mesenchymal cell types in the bone marrow. To achieve this, we employed whole-slide CODEX multiplexed imaging of 12 specimens, 8 of which were included in our transcriptomic atlas (Figure 4A). Guided by our scRNA-seq data, we designed and validated a 54 marker panel to interrogate the bone marrow microenvironment (Figure 4B, Supplemental Figure 4A, Supplemental Table S4-S6). We included markers such as SPINK2, which was expressed most highly in scRNA-seq defined primitive HSCs (Supplemental Table S2), to increase the resolution of primitive HSPCs in addition to traditional markers such as CD34 and CD38. Whole-slide CODEX images were acquired and subsequently segmented using Mesmer^45^ to identify single cells. Cells were then annotated by a combination of iterative clustering and, in rare circumstances, cluster annotation refinement by manual gating. Importantly, after each round of clustering, cell labels were overlaid on the fluorescent image and clusters were visually inspected and corrected if necessary (Supplemental Figure S4A). In sum, we computationally annotated 803,132 cells (91.6% of segmented objects) (Figure 4C). We identified 32 cell types across 12 samples, including cells as rare as immunophenotypic HSCs (Lin-CD34+ CD38-CD45RA-CD90+) and Schwann cells (PLP1+ CD271+) (Figure 4C). Hematopoietic cells were annotated by canonical surface marker expression (Supplemental Table S4). Furthermore, we subcategorized the non-hematopoietic cells into AECs, SECs, VSMCs, endosteal cells, Adipo-MSCs, and THY1+ MSCs. We also detected a population not captured in our scRNA-Seq due to tissue processing and abundance – Schwann cells. Notably, despite some tissue processing artifacts, such as frequent loss of bone tissue, we were still able to identify para-trabecular cells with non-hematopoietic expression profiles that we termed “endosteal”. We detected a cluster corresponding to adipocytes but also tissue processing artifacts, which we termed adipocyte/artifact. None of Osteo-MSCs, Osteo-Fibro MSCs, or Fibro-MSCs were detected computationally, suggesting these cells are tied more closely to bone and are thus lost in the tissue processing. We detected rare instances of these cells (Supplemental Figure S4B-C), with osteo-MSCs and osteoblasts being detected on the hematopoietic-bone interface, but Fibro-MSCs being detected largely in the middle of the bone region. Therefore, we focused on Adipo and THY1+ MSC in our CODEX analysis. For brevity, “MSCs” will hereafter refer to Adipo and THY1+ MSCs collectively, unless otherwise specified. Interestingly, we also found that all macrophages observed were M2-like as defined by CD68/CD163 co-expression, and that megakaryocytes could be split into GATA1-positive and GATA1-negative subtypes (Figure 4C, Supplemental Figure 4D). Broadly speaking, the cell type distribution was very similar between patients (Figure 4D, Supplemental Table S1). These annotations comprise a digitized record of nearly every cell across twelve bone marrow samples, and we can visualize the cell type organization on a single-cell level (Figure 4E, Supplemental Figure 4A). Collectively, this annotated reference serves as a comprehensive map of the cellular topology of hematopoiesis in the context of its microenvironment and demonstrates the complementarity of these two atlases to understand the cellular anatomy of human bone marrow.

**Figure 4.**
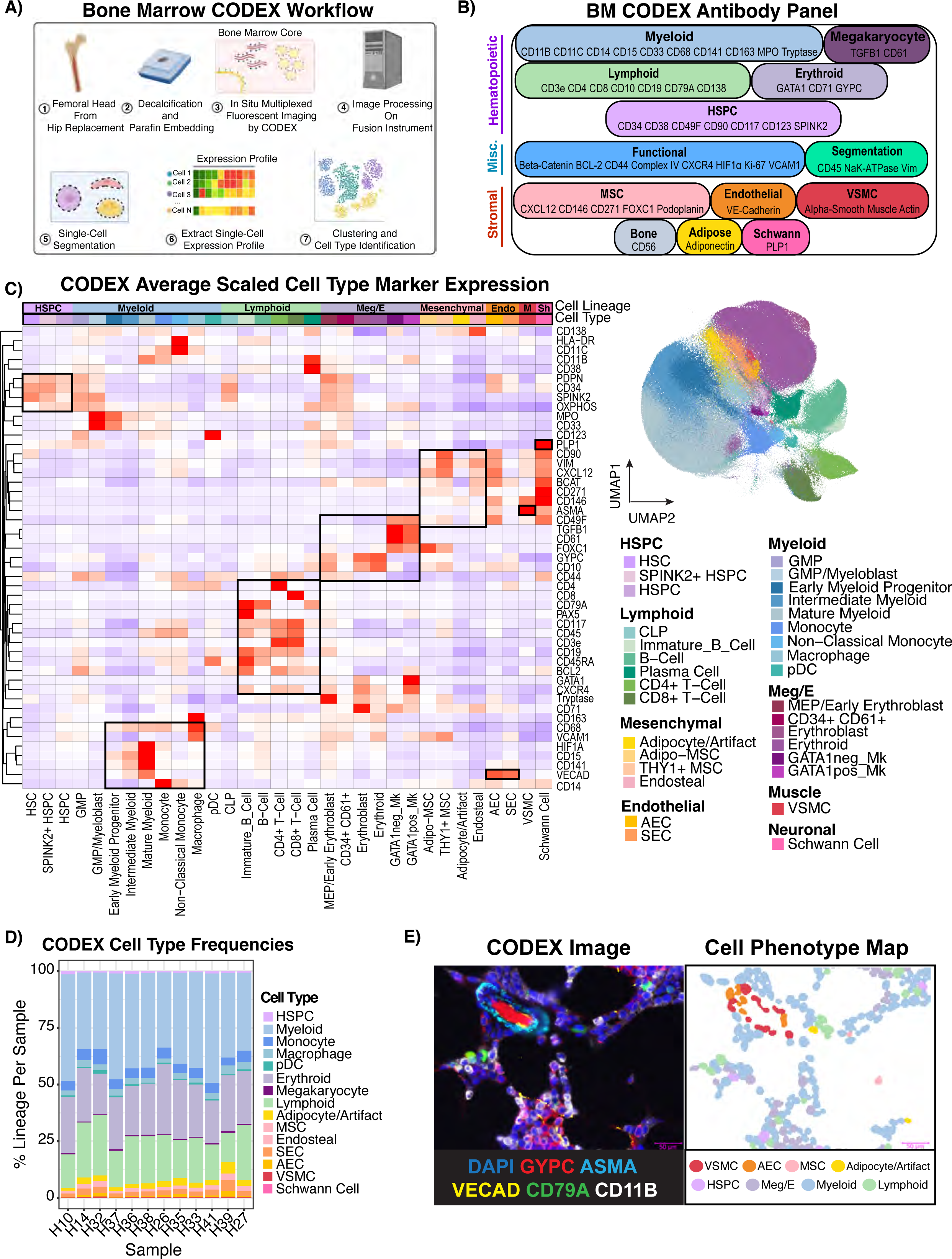
54-plex CODEX imaging reveals the spatial cellular topography of human bone marrow. **A)** Schematic depicting the CODEX experimental and computational workflow leading to cell type identification. **B)** Diagram showing the 54-plex CODEX panel (53 antibodies + DAPI) split by target cell population. **C)** Heatmap showing average centered-log-odds ratio normalized expression per cell type scaled and clustered by protein showing the marker protein expression is consistent with literature knowledge (Left). Cell types were annotated by unsupervised clustering and manual gating as described in Materials and Methods. Boxes were manually drawn to highlight coordinate marker expression. UMAP showing the 803,131 single cells in the CODEX atlas from 12 individuals colored by cell type (Right). Adipocytes were labeled with /Artifact because this population contained a mixture of true adipocytes and artifactual staining. HSPC – Hematopoietic Stem and Progenitor Cell. Meg/E-Megakaryocyte/Erythroid, Endo-Endothelial, M-Vascular Smooth Muscle, Sh-Schwann Cells. **D)** Stacked bar plot showing cell type frequencies normalized by total cells per sample. **E)** CODEX image (left) is paired with the cell phenotype map (CPM,right). An arteriolar structure and hematopoietic cells are shown using selected relevant fluorescent markers, which is juxtaposed with the same image with the cellular segmentation masks colored by cell type showing how CODEX allows single-cell mapping of the bone marrow microenvironment.

### Early Myeloid Progenitors and GMPs Consistently Localize to a Relatively Hyperoxygenated Arterio-Endosteal Niche

While multiple niches such as endosteal, perivascular, and arteriolar have been proposed to organize hematopoiesis in extensive and often controversial literature^2,5,8,32^, we sought to utilize an unbiased, statistically rigorous approach to answer the question of how hematopoietic and mesenchymal cells organize into cellular communities. Indeed, even with interactive visualization of the CODEX images, we noticed discrete patterns of cellular organization, such as peri-endosteal localization of MPO^hi^ early myeloid progenitors (EMPs), or erythroblastic islands composed of CD163+ macrophages and erythroid precursors (Figure 5A). We therefore performed unsupervised neighborhood analysis^46^ and identified 15 cellular neighborhoods (CNs) which we manually annotated based on their relative enrichment of cell types (Figure 5B). CNs with similar major cell types and thus same annotations were manually combined. Given the challenge of bone detachment during the imaging, we measured neighborhood assignments with respect to manually annotated bone locations, confirming that neighborhoods enriched for “endosteal” cells were indeed close to bone (Supplemental Figure S5A). We found multiple mixed lineage neighborhoods with all three hematopoietic lineages (CN7, CN8, i.e. Erythroid/Myeloid/Lymphoid) (Figure 5B). We also found expected neighborhoods such as the “Erythroid” neighborhood (CN13, CN15) consistent with erythroblastic islands, and also novel neighborhoods, such as one neighborhood of peri-arteriolar lymphoid cells (Figure 5B, Supplemental Figure S5B). Consistent with recent reports of their random distribution throughout the marrow^47^, HSPCs did not have a strong preference for one particular neighborhood, and this was true for immunophenotypic HSC, SPINK2+ HSPC, and all other HSPC. HSPCs were found most strongly in mixed erythroid/myeloid/lymphoid neighborhoods (CN7, CN8), and to a lesser degree in Myeloid/Lymphoid (CN5) and Early Myeloid/Arteriolar (CN4) neighborhoods (Figure 5B). We also recognized two neighborhoods (CN4,CN6) enriched for GMP/myeloblasts and early myeloid precursors heavily enriched along the endosteal surface and around arterioles (Figure 5B-C). This was also consistent with the endosteal signaling module we identified using CellChat which had contribution from AECs and VSMCs. We therefore termed them Early Myeloid/Arteriolar(CN4) and Early Myeloid/Endosteal (CN6), collectively Early Myeloid, based on the enrichment scores for AECs and endosteal cells respectively (Figure 5B-C). Notably, this was distinct from the peri-arteriolar lymphoid niche, demonstrating heterogeneity even within arteriolar niches (Figure 5B, Supplemental Figure S5B). All of the arteriolar neighborhoods (CN1-CN4), including Early Myeloid/Arteriolar(CN4), were enriched for Adipo- and THY1+ MSCs (Figure 5B). Importantly, this analysis also shows that the concept of the peri-arteriolar and endosteal niches holds up to unbiased, statistically-based measurement of spatial association, but suggests that in line with recent studies using mouse models^47^, most human HSPCs do not preferentially occupy an endosteal or peri-arteriolar niche.

**Figure 5.**
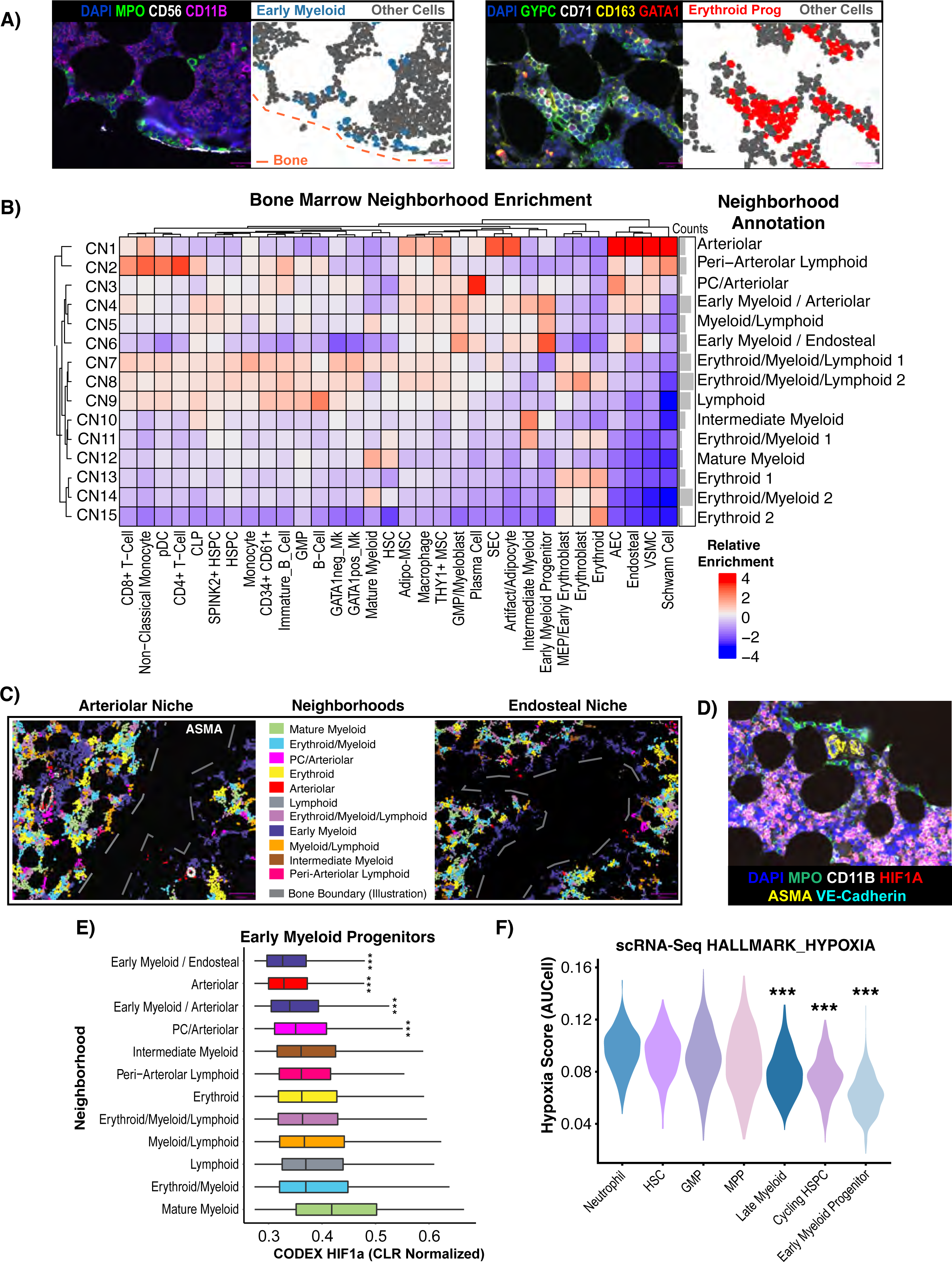
Neighborhood analysis reveals an oxygen-rich arteriolar/endosteal niche for GMP/early myeloid progenitor cells. **A)** CODEX images juxtaposed with cell phenotype maps (CPMs) showing endosteal localization pattern of MPO+ early myeloid progenitors (CPM of EMPs) (left) and erythroblastic islands with CD163+ macrophages and GYPC+ CD71+ erythroid progenitors (CPM of EPs) (right). All other cells were colored grey. **B**) Neighborhood analysis performed by clustering the cell type frequency vectors of the 10 nearest neighbors of each cell, and a heatmap was generated showing the relative enrichment of each cell type normalized by neighborhood. Neighborhood names were manually assigned based on the relative enrichment of the cell types present. Downstream of this, neighborhoods with the same manual annotation (e.g. Erythroid 1 and Erythroid 2) are combined. **C**) Cell phenotype masks colored by neighborhood membership were plotted alongside ASMA fluorescent signal (left only). Arteriolar and endosteal niches for early myeloid progenitors were made the same color for the purpose of this visualization. Bone boundaries were manually annotated due to the bone typically detaching from the slide during the antigen retrieval process. **D**) Fluorescent CODEX image showing the endosteal MPO+ early myeloid progenitors not expressing HIF1a, in contrast to more central, mature myeloid elements. **E**) Boxplots showing the centered log ratio (CLR)-normalized HIF1a expression levels in early myeloid progenitors split by neighborhood membership, showing that HIF1a levels vary between neighborhoods in addition to cell type. **F**) Violin plots showing that the populations identified by scRNA-Seq which are analogous to early myeloid progenitors in CODEX data also have low GSEA Hallmark hypoxia gene set scores calculated by AUCell in the scRNA-Seq data.

The importance of the endosteal niche has been previously reported in both healthy and malignant hematopoiesis^8,48^. We therefore sought to further investigate interactions within this niche using both our CODEX and scRNA-Seq atlases. First, we considered that the dual peri-arteriolar and peri-endosteal localization of early myeloid progenitors could be related. We found that arteriolar cells were much more frequently found near the trabecular bone (Figure 5B, Supplemental Figure 5A), suggesting that the endosteal niche may actually combine with the arteriolar niche. Furthermore, we found that HIF1a levels were nearly undetectable in early myeloid progenitors and GMPs, suggesting that these cells are not experiencing hypoxia (Figure 5D), correlated with their localization near oxygen-supplying vessels. In contrast, more mature myeloid cells had extremely high levels of HIF1a (Supplemental Figure S5C). We found that early myeloid precursors found in non-arteriolar neighborhoods had higher levels of HIF1a, demonstrating this is likely to reflect spatial patterns of hypoxia rather than cell type difference alone (Figure 5E). We then employed AUCell^49^ analysis of our scRNA-Seq atlas to interrogate whether a hypoxic gene signature was differentially expressed at different stages of myelopoiesis. In support of the CODEX findings, we found that early myeloid progenitors have the lowest hypoxia signature score (Figure 5F). Given that the bone marrow is broadly a hypoxic environment, this finding suggests that a relatively oxygenated niche is critical for early myelopoiesis. Furthermore, this suggests that the arteriolar and endosteal niches are not, as is often discussed in the bone marrow niche literature^2,5^, two entirely discrete niches and that there is significant proximity and likely crosstalk between these two parts of the bone marrow microenvironment.

### Imaging-based Structural Analysis Reveals Peri-Adipocytic Localization HSPCs

We next considered that certain key microenvironmental structures such as bone, adipocytes, stroma, sinusoids, arterioles, and macrophages have complicated 3D architecture which may not be captured by classical cell segmentation approaches. We employed manual annotation and thresholding to confidently annotate six major structures – adipocytes, arterioles, bone, macrophages, sinusoids, and CXCL12+ stroma (Supplemental Figure S6A). We implemented point pattern analysis (See STAR methods) to analyze the distance between bone marrow cell types and neighborhoods to each of these newly annotated structures^50^. Each structure, neighborhood, and cell type were ranked based on their proximity to each microenvironmental structure. We found that of all the measured structures, the structure closest to bone were the arterioles (Figure 6A). This supports our observation that the endosteal niche was arteriole-rich (Figure 6B). In general, we found that the central niche, as defined by being far from the bone, was characterized by sinusoids, adipocytes, stroma, and macrophages, although adipocytes and sinusoids were further away than stroma or macrophages (Figure 6A-B). Additionally, structural analysis confirmed that the previously mentioned erythroid niche corresponded to erythroblastic islands, as this niche was very proximal to macrophages (Figure 6C). The early myeloid neighborhoods were both close to arterioles and bone, providing further evidence of the arterio-endosteal EMP niche (Figure 6C).

**Figure 6.**
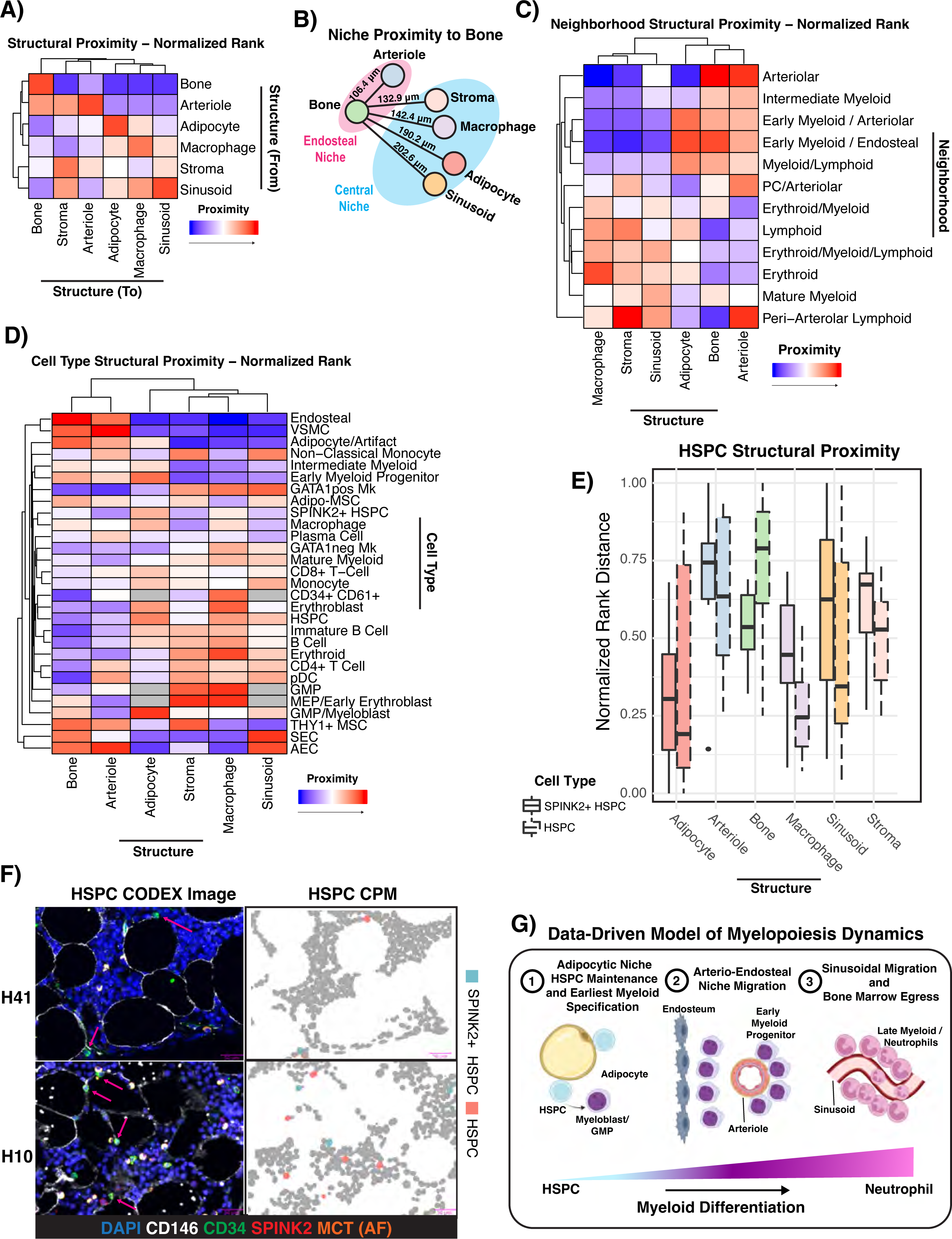
Comprehensive structural analysis of bone marrow microenvironment uncovers adipocytic localization of Lin-HSPCs. **A)** Hierarchically clustered heatmap showing the normalized rank proximity from each microenvironmental structure to each structure. Data is from all 12 samples. Each structure was ranked by its proximity to each other structure in each sample, and the rank was normalized such that a value of 0 means most proximal and 1 means least proximal. The color scale was adjusted such that 0 is red and 1 is blue. **B)** Data-driven illustration of structural proximity in normal bone marrow. Edge lengths are scaled to the median normalized rank proximity (where lower numbers mean more proximal) of each structure, and the physical median distance across 12 samples is labeled above each line. **C-D)** Hierarchically clustered heatmap showing the normalized rank proximity of each neighborhood and cell type to each type of microenvironmental structure. Data is from all 12 samples, except when a cell type or neighborhood was too rare to rank in a sample. Each cell type or neighborhood was ranked by its proximity to each structure in each sample, and the rank was normalized such that a value of 0 means most proximal and 1 means least proximal. The color scale was adjusted such that 0 is red and 1 is blue. Grey color means that there weren’t enough cells (<10 in at least half of samples) in the annotated region to include in the analysis for that cell type. Immunophenotypic HSCs, CLPs, and Schwann cells were excluded completely from the cell type analysis by this criterion. **E)** Boxplot showing the normalized rank proximity of Lin-HSPCs (dashed outline) and Lin-SPINK2+ HSPCs (solid outline) to each microenvironmental structure. **F)** Representative CODEX images and associated CPMs from two samples showing the peri-adipocytic localization of both HSPCs and SPINK2+ HSPCs. **G)** Cartoon schematic showing the revised model of human myelopoiesis, where adipocyte-adjacent HSPCs give rise to the earliest myeloid committed cells, which migrate to mature in an arterio-endosteal niche, and finish their maturation and egress in the sinusoidal/central niche.

Our results also validated our cell type annotation, as the expected cell types co-localized with the expected structures such as MSCs-stroma, macrophages-macrophages, endosteal-bone, SEC-sinusoids, etc. (Figure 6D, Supplemental Figure S6B). Interestingly, we found that all MSCs were close to the arterio-endosteal niche, but Adipo-MSCs were found near sinusoids while THY1+ MSCs were not (Figure 6D). This may suggest that Adipo-MSCs are more analogous to previously described HSC-supportive peri-sinusoidal MSCs^37,51^.

We also discovered spatial patterns not apparent in the previous cell segmentation-based analysis. HSPCs and more primitive SPINK2+ HSPCs were both found to be highly proximal to adipocytes (p=0.0198 and p=1.77E-6 respectively using permutation testing, Figure 6D-E). Manual inspection of CODEX images and CPMs revealed that both Lin-SPINK2+ CD34+ and Lin-CD34+ cells were in frequent contact with adipocytes (Figure 6F, Supplemental Table S7). This extends and contextualizes previous findings^52^ with respect to the entire bone marrow microenvironment, and demonstrates that Lin-SPINK2 +/- CD34+ HSPCs have the strongest spatial association with adipocytes of all microenvironmental structures measured. Interestingly, this finding extended to GMP/Myeloblast cells as well as early myeloid progenitors, although these cells were significantly close to bone while HSPCs were not (Supplemental Table S7). HSPCs, but not more primitive SPINK2+ HSPCs, were also found to be close to macrophages as well (p=5.23E-11and n.s. respectively, Figure 6E). Notably, mature myeloid cells, by contrast, were closest to sinusoids and not arterioles or bone, suggesting that maturation occurs in a central, peri-sinusoidal niche (p=6.71E-68, Figure 6D). Collectively, this implies spatial restriction of myeloid development where the earliest myeloid progenitors are specified in an adipocytic niche from HSPCs, migrate to a relatively hyperoxygenated arterio-endosteal early myeloid niche, and finally migrate to mature near sinusoids they may utilize to egress from the marrow and into the bloodstream. This analysis enabled us to make a revised schematic of the bone marrow niche organization, which should motivate future functional studies which aim to interrogate the biological mechanisms underlying this spatial organization (Figure 6G).

### Unsupervised Reference Mapping Using Healthy NBM CODEX Atlas Reveals AML Stromal Expansion and Novel AML-MSC Enriched Neighborhoods

Understanding the heterogeneity of the bone marrow microenvironment has significant implications for disease states as well as healthy hematopoiesis. We therefore sought to determine whether our healthy atlas could be leveraged to study the bone marrow microenvironment in the context of malignancy. Acute Myeloid Leukemia (AML) is a cancer which evolves from accumulation of mutations in HSPCs^53^ and has been described previously to interact with the endosteal niche to achieve chemoresistance^48^. We used our bone marrow CODEX panel to explore tumor evolution and microenvironmental changes using our healthy atlas as a reference. We profiled three diagnostic and two paired post-therapy bone marrow biopsies from AML patients (Supplemental Table S1). The patients were treated with venetoclax and a hypomethylating agent. We also included similarly processed negative lymphoma staging bone marrow biopsies (NSM) as controls. NSM samples were chosen as the control to account for differences between sample site (femoral head vs. iliac crest), sample processing, and generalized stress associated with having cancer. We selected AML patients with a detected *NPM1* mutation (*NPM1c* W288*fs) at diagnosis and after therapy due to availability of a mutant specific antibody^54,55^ so that we could readily identify putative leukemic blasts. We employed reciprocal principal component analysis (RPCA) to classify cells in the AML and NSM samples to their nearest counterpart in our healthy atlas (Figure 7A). Using this approach, we found that as expected, there was a significant increase in the fraction of cells of the myeloid compartment in diagnostic AML compared to NSM (p<2.2E-16,Figure 7B). This demonstrates the flexibility of the atlas to rapidly map and label hundreds of thousands of cells in both healthy and diseased contexts. Projecting these cell labels onto the segmentation masks clearly illustrates grossly appreciable architectural changes in NSM, diagnostic AML, and post-therapy AML – with a near total loss of certain mesenchymal marrow elements such as adipocytes in the leukemic setting, with incomplete recovery 30 days post-therapy in the presence of residual blasts (Figure 7C). A more detailed look at the cellular landscape of AML also revealed marked stromal expansion, with 2-3 fold higher relative frequency of Adipo and THY1+ MSCs in AML samples compared to NSM (Figure 7D). Next, we aimed to define leukemic blasts in these tissues to help determine whether these blasts preferentially associated with MSCs or other cell types. We observed that the NPM1 mutant antibody had low-level of binding to cytoplasmic elements in more mature myeloid cells which expressed CD141 and plasma cells marked by CD38 and CD79A (Supplemental Figure S7A-B). To circumvent this issue, we trained a classifier to identify true *NPM1* mutant cells based on marker expression patterns which were exclusive to the AML samples. Furthermore, we filtered out any leukemic cells which mapped to incompatible or very rare cell types. Our classified blast percentages correlated with the *NPM1c* variant allele frequencies (Supplemental Figure S7C). Using this approach, we were able to label and visualize leukemic populations including a GATA1+ NPM1c+ double positive population in the post-therapy sample of AML3 and identify individual *NPM1* mutant MRD cells (Figure 7E). We did observe a ∼2-fold increase in GATA1 expression in *NPM1* mutant blasts across both post-therapy samples, suggesting that these MRD cells exhibit a high degree of lineage plasticity (p<2.2E-16, Figure 7E). We next profiled spatial patterns between cell types in leukemia and NSM samples. We used unsupervised neighborhood analysis to identify 15 cellular neighborhoods, of which three were enriched for *NPM1* mutant cells – CN2, CN13, and CN15 (Figure 7F). All three of these leukemic neighborhoods were enriched for MSCs, which we observed visually as well (Supplemental Figure S7D). CN2 and CN13 were enriched for VSMCs, endosteal cells, monocyte/DCs, and B cells as well. Importantly, all three of these neighborhoods were specific to the AML samples, implying an organizational pattern that is unique to leukemia (Figure 7G). CN2 and CN13 had similarities to CN7, a non-leukemic neighborhood found roughly equally in all three sample types which corresponded to the previously described early myeloid progenitor niche with enriched MSCs, endosteal cells, and early myeloid progenitors. Notably, unlike CN7, leukemic neighborhoods were not measurably close to bone, suggesting that another factor such as MSCs may be the driver of leukemic neighborhood localization (Supplemental Figure S7E). We also noticed that the leukemia-specific neighborhoods were preserved in the post-therapy setting. Both diagnostic and MRD *NPM1* mutant blasts had low HIF1a levels consistent with GMPs and early myeloid cells (Supplemental Figure S7F), and for the two patients with paired samples, MRD cells had slightly decreased BCL2 and increased mitochondrial Complex IV after therapy (p < 2.2 E-16 for both, Supplemental Figure 7G-H). This could reflect decreased reliance of these MRD cells on the BCL2 pathway and compensatory increase in mitochondrial mass to circumvent venetoclax-mediated suppression of BCL2 and oxidative phosphorylation^56,57^. These data provide direct *in vivo* evidence of AML-MSC spatial interactions. Taken together, these results highlight the broad utility of our healthy bone marrow CODEX reference, demonstrate the power of high throughput computational image analysis to illuminate spatial biology in bone marrow disease states, and provide further rationale to study stromal-leukemic cell interactions in AML.

**Figure 7.**
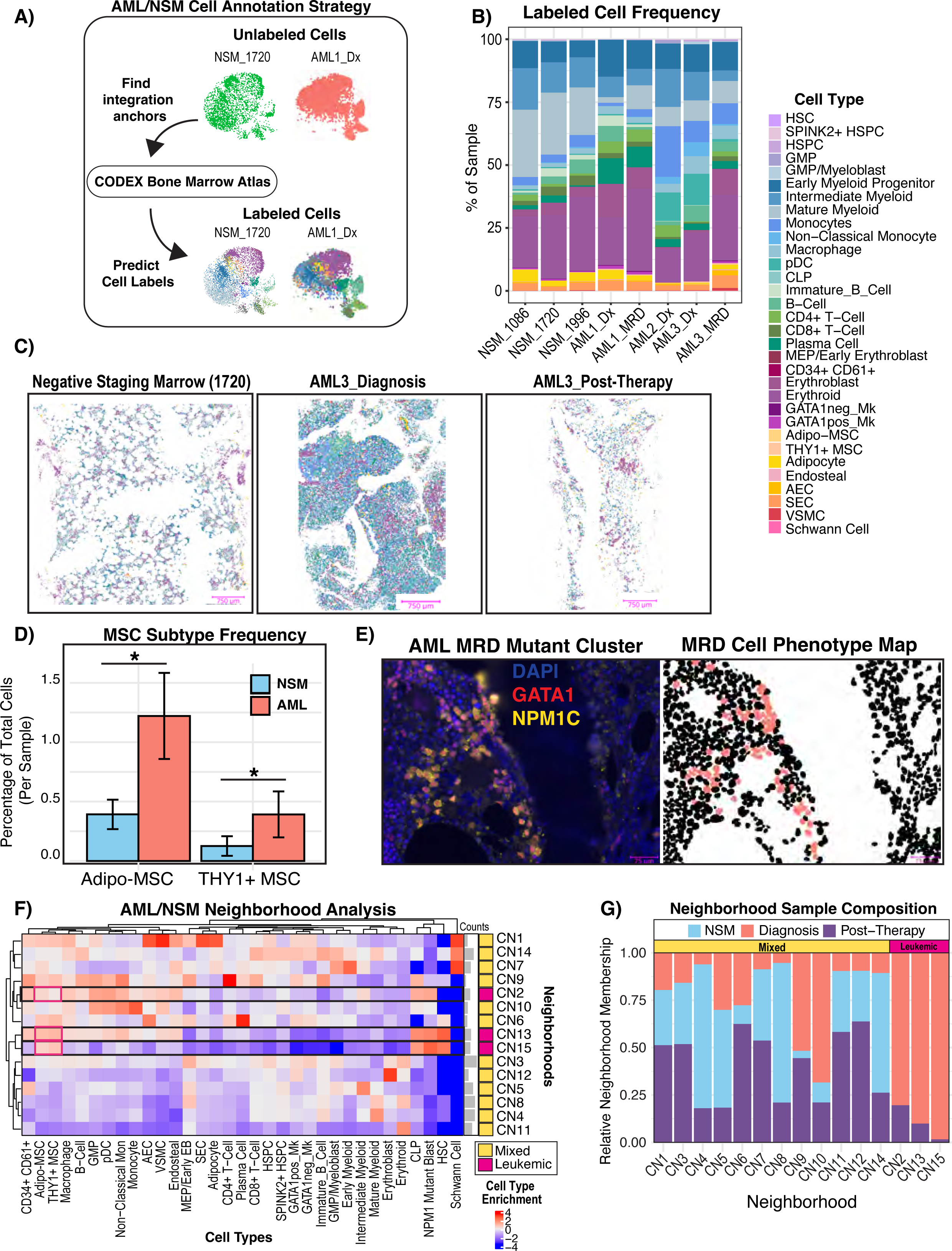
Unsupervised single-cell mapping of AML reveals stromal expansion and MSC-enriched AML-specific neighborhoods. **A**) Schematic showing our unsupervised label transfer computational approach. Reciprocal PCA (RPCA) integration was used to predict cell labels in NSM and AML samples based on their proteomic similarity to the annotated normal atlas presented in Figure 4. **B)** Bar plot showing the cell type frequencies in each sample. AML1_Dx: n= 9,334 cells, AML1_MRD: n= 68808 cells, AML2_Dx: n-63412 cells, AML3_Dx: n= 37063 cells, AML3_MRD: n = 17,035 cells, NSM_1086: n =14,097 cells, NSM_1720: n = 118,643 cells, NSM_1996: n= 32,560 cells **C)** Cell phenotype maps (CPMs) shown for representative images of each sample. Masks are colored by cell type and share a legend with panel B. **D)** Bar plot showing per sample frequency of MSCs in AML (both diagnostic and MRD) vs NSM. MSC frequency was calculated as the proportion of cells in each sample which had the predicted label “Adipo-MSC” or “THY1+ MSC”. The distribution of MSC subtype frequencies in AML sample was compared to that of NSM samples using a two-sided t-test. **E)** CODEX image showing clustering of rare residual *NPM1* mutant blasts which also stained positive for GATA1, juxtaposed with the CPM showing these mapped MRD cell (salmon color) locations. The blank region extending from top to bottom separating the two cellular regions is bone. **F)** Heatmap showing the neighborhood cell enrichments (artifacts removed, AML: n=175365, NSM: n = 152198). Cell type enrichment (fold change) was calculated as previously described^46^ and scaled by cell type frequency (by column) to highlight which neighborhoods most of each cell type was found in. **G)** Stacked bar plot showing the relative frequency of each sample type in each neighborhood. Frequencies were normalized by total cell number per sample type (AML_Diagnosis, AML_Post-Therapy, NSM) so that total cell count differences between the sample groups do not skew the results. The mixed neighborhoods were defined as any neighborhood which had cells from both AML and NSM samples, whereas AML-specific neighborhoods were defined by only being found in diagnostic or post-therapy AML samples.

## Discussion

The non-hematopoietic human bone marrow microenvironment remains largely uncharacterized. Here, we provide an atlas with two complementary modalities which collectively represent the first comprehensive healthy human bone marrow reference which profiles the phenotype and biogeography of both hematopoietic and non-hematopoietic cell types revealing the extent of cellular diversity and spatial organization.

With our novel human bone marrow processing protocol, we identified six mesenchymal subsets, two endothelial cell subsets, and one vascular smooth muscle population. Of these, Fibro-MSCs were computationally and functionally shown to be the most in accordance with human mesenchymal stem cells described previously^26^. Our 82,743 single-cell transcriptomic atlas, comprising 53,417 hematopoietic and 29,325 non-hematopoietic cells, is the largest comprehensive atlas of healthy human bone marrow. Our atlas can be used to contextualize diverse mesenchymal cells in existing and future bone marrow scRNA-Seq datasets, exemplified by the usage to explain relative homogeneity in bone marrow aspirate MSCs^19^ or to elucidate the differential composition of the fetal bone marrow niche as compared to adult marrow^31^. It should be noted, however, that our atlas may have limited resolution in deconvoluting the cell types absent in typical older adult bone marrow, motivating the application of this experimental approach to generate analogous pediatric and fetal references in the future. These data have important implications for future studies of the human bone marrow microenvironment and demonstrate the importance of isolation method and non-hematopoietic cell enrichment.

We applied the insights gained from our transcriptomic analysis to devise a 54-marker CODEX bone marrow panel which elucidated the spatial relationships between the bone marrow microenvironment cell types. Our 803,131 cell healthy spatial proteomic dataset is the first human atlas which spatially profiles discrete stages of erythropoiesis, lymphopoiesis, and myelopoiesis in the context of all major microenvironmental structures. This resource will be critical for future study of bone marrow organization, especially as spatial imaging analysis methods grow more sophisticated. This work builds upon decades of literature debating the importance of a perivascular or endosteal niche for HSPC maintenance^2,5,51^. These studies are often limited, however, by utilization of arbitrary metrics for spatial proximity, or bias in studying one element of the microenvironment at a time. In this study, we perform unbiased, comprehensive analysis of the cell types and microenvironmental structures found in human bone marrow which enabled new insights. For instance, recent murine models^47,58^ support a random distribution for HSPCs, but our atlas reveals that a cell type not assessed explicitly in those studies, adipocytes, are the structure that human HSPCs are closest to. This finding is of particular importance as adipocytic density increases as humans age and local adipocytic density is markedly reduced in the context of acute leukemia. However, bone marrow adipogenesis has been proposed as a negative regulator of hematopoiesis in mice^59^, and so whether this interaction represents a hematopoiesis-supportive or suppressive interaction in humans warrants further investigation. Either way, our data did not support an endosteal or perivascular preference for human HSPCs. Importantly, this data does not mean that sinusoid-HSC communication is unimportant, only that human HSPCs aren’t closer to sinusoids than expected by random chance. We also find significant evidence of spatial restriction of myelopoiesis, whereby myeloid progenitors migrate from adipocytic niches at the HSPC stage, to relatively oxygenated arterio-endosteal niches in the myeloblast/promyelocyte stage, and then to the sinusoids as they mature and egress from the bone marrow. Our data demonstrate the heterogeneity of peri-vascular niches in the bone marrow, quantitatively connect the arterial and endosteal niches, and show that sinusoids are just one component of a complex multicellular spatial organization. This quantitative delineation of the bone marrow niche’s constituents will be crucial to the continued efforts to model the human bone marrow, such as in recent organoid or microfluidic chip systems^60–62^, particularly as current iterations of these models typically contain only a subset of the bone marrow non-hematopoietic cells.

We further demonstrated the utility of this atlas by applying it to the goal of contextualizing disease. We found that *NPM1* mutant blasts in both diagnostic and post-therapy settings are found in MSC-enriched neighborhoods. We show that it is both feasible and informative to use CODEX to map blast-stromal interactions to study the bone marrow microenvironment, and this work provides further rationale to study stromal-blast interactions in the setting of AML. Furthermore, our ability to detect and map leukemic MRD cells highlights the novel translational potential of bone marrow CODEX to detect and find prognostic spatial biomarkers of residual disease, which is known to be associated with increased rate of relapse and poor outcome in AML^63^, and particularly in *NPM1* and *FLT3* mutant patients^64^. Collectively, these results highlights the potential of our comprehensive spatial atlas to uncover novel biology in both healthy and disease settings.

Taken together, this study provides the first transcriptionally and spatially resolved atlas of the human bone marrow microenvironment. In a highly contentious field, there is a growing need for more standardized, quantitative, and unbiased metrics to reconcile the cumulative body of research on the bone marrow microenvironment over the past few decades. Here, we took such a systems biology approach to reveal the cellular heterogeneity and biogeography of the bone marrow. We believe this atlas will serve as a critical resource for future studies of the human bone marrow niche.

### Limitations of the Study

It should be noted there are some limitations with the patient cohort, imaging analysis, and inherent limitations of the imaging methodology.

Our patient cohort comprises a representative sample of twelve individuals aged roughly 50-70 years, approximating a relatively “old” microenvironment. However, the human bone marrow changes drastically from the fetal, post-natal, infant, pediatric, young adult, and older adult stages. Therefore, significant future work remains to determine the temporo-spatial landscape of hematopoiesis in humans.

Additionally, image analysis in the context of mapping complex cellular topologies, so-called “neighborhoods”, is in its infancy. Tools do not currently exist which combine prior knowledge regarding secondary or tertiary biological structures (e.g. arterioles or lymphoid follicles) along with cellular data for quantitative neighborhood analysis, which could help resolve interaction between higher-order structures and individual cells.

Furthermore, 2D imaging is inherently noisy, limited to distances which can be measured by observable analytes within a 2D plane within a complex 3D structure. This is particularly problematic for very rare cell types like hematopoietic stem cells, which could have supportive structures directly above or below the captured plane but would not have a spatial association captured. As 3D multiplexed imaging and corresponding analytical approaches develop, we will gain a more comprehensive quantitative view of the human bone marrow.

## Methods

### Human clinical sample acquisition

Bone marrow samples were obtained from patients undergoing total hip arthroplasty surgery. We reasoned that osteoarthritis patients undergoing total hip arthroplasty surgeries, where the femoral bone is removed and discarded as surgical waste, would be a good source of hematologically healthy bone marrow. We considered that these samples were representative of normal hematopoiesis due to two major factors: first, studies have reported that 75-90% of individuals over 75 years of age will develop some degree of osteoarthritis^65^, making the condition within the range of aging-associated changes; and second, osteoarthritis is a peri-articular disease not associated with hematologic complications^66^. We only sampled at the distal site to the articular cartilage and only collected morphologically normal trabecular bone and associated bone marrow. For sequenced femoral heads, the age range was 52-74 years, with a median of 65+/- 8 years. Patients filled out a questionnaire explaining whether they had certain common medical conditions which are tabulated in the clinical metadata table. All patient samples were deemed as medical waste and IRB exempt by the University of Pennsylvania Institutional Review Board. Only non-identifiable patient information is collected for each sample by the clinical staff. Researchers were never given access to medical records. AML and NSM samples were obtained from the Hospital of the University of Pennsylvania’s Pathology sample repository.

### Femoral head cell isolation

Femoral heads obtained from total hip arthroplasty surgeries were cut coronally in 4-6 mm thick slices by the Anatomical Pathology Lab at Penn Presbyterian Medical Center. Tissue samples were transported on ice in AlphaMEM (Gibco) with 3% FBS (HyClone), 1% penicillin-streptomycin (Gibco), 0.02 mM L-ascorbic acid (Sigma) and 0.3 mM L-Glutamine (Gibco). Samples were placed in a petri dish and cut with surgical wire cutters into approximately 1 mm cubed pieces, weighed and transferred to a 15 mL centrifuge tube. For CODEX, a dental biopsy drill was used to harvest cylindrical samples, cut to about 4 mm thick, and fixed in 4% paraformaldehyde (PFA) (EMS). For scRNA-Seq and cell sorting, 1 mm of minced tissue was digested in filtered 4 mg/mL dispase II (Sigma), and 2 mg/mL collagenase I (Worthington Biochem) in PBS for one hour on a rocker at 37 °C. The cell suspension was then filtered through a 100 micron cell strainer (Fisher) and remaining tissue was washed five times in culture medium (AlphaMEM with 15 % FBS, 1% P/S, 0.1 mM LAA, 1 mM L-glutamine) and strained. Total collected suspension was pelleted at 300 RCF for 10 mins and resuspended in MesenCult Proliferation medium (STEMCELL, Catalog # 05411), and immediately taken for immunomagnetic population isolation (see “Single-Cell RNA sequencing” or “Mesenchymal cell fluorescence assisted cell sorting”).

### Micro-CT BV/TV determination

Bone tissue cores were scanned with a micro–computed tomography (micro-CT) 35 scanner (Scanco Medical AG, Switzerland) at a 6-μm isotropic voxel size. All images were analyzed with the Scano scanner software and a region 4200 microns in diameter and 900 microns tall was selected and smoothed with a Gaussian filter (sigma = 1.2, support = 2.0) prior to binarization with a minimum threshold of 365 and bone volume to total volume (BV/TV) calculated.

### Single-Cell RNA sequencing

Digested tissue samples were subjected to immunomagnetic RBC depletion reagent (STEMCELL #18170). 20uL of RBC depleted cells were put on ice, while 67% of the remaining cells were used for immunomagnetic CD45 depletion using EasySep Human CD45 Depletion Kit II (STEMCELL # 17898) to isolate non-hematopoietic cells in an unbiased fashion, while 33% of cells were subjected to immunomagnetic CD34 enrichment using EasySep Human CD34 Positive Selection Kit II (STEMCELL #17856). Following enrichment, each of the three cell fractions (RBC depleted, CD45 depleted, CD34 enriched) were resuspended at a concentration of 1000 cells / uL in PBS + 0.04% BSA. Cells were pooled and loaded into the 10x Genomics Chip G (#1000127). We used Chromium Next GEM Single Cell 3’ GEM, Library & Gel Bead Kit v3.1 (10x Genomics, #1000121) and followed the protocol as provided by 10x Genomics. Sequencing was performed at the High Throughput Sequencing Core at CHOP using a NovaSeq 6000 system. Sequencing parameters were set according to 10x Genomics recommendations.

### scRNA-Seq data processing

scRNA-seq data was demultiplexed, aligned, and count matrices generated using Cell Ranger v6.1.2 aligning to GRCh38 with intronic regions included. Cell Ranger matrices were directly imported into Seurat v4. We used DoubletFinder^67^ to remove doublets on a per-sample basis. Cells with 100-10,000 unique genes expressed and less than 10% mitochondrial reads were retained for the downstream analysis. Samples were combined without computational integration and subjected to dimension reduction and clustering. Cells were annotated by marker gene expression, and likely artifactual cells (only found in one sample or likely residual doublets) were removed from analysis using VisCello for interactive visualization and artifact selection and removal^68^. Mesenchymal and endothelial cell clusters were computationally isolated for further analysis. AUCell^49^ was used for pathway analysis with the GSEA human Hallmark gene set (http://www.gsea-msigdb.org/gsea/msigdb/collections.jsp). Further details are provided in the accompanying source code.

### Cellular interaction analysis

CellChat was used for cellular interaction analysis using the CellChatDB database. As described in the original manuscript^33^, scRNA-Seq values were projected onto a protein-protein interaction network to account for typical dropout issues. The significant cell-cell interactions were determined using CellChat default parameters, with population size accounted for and “trimean” used for calculating mean gene expression. Pathway annotations were taken from the original manuscript. For the Non-negative Matrix Factorization (NMF) signaling pattern analysis, K was selected by taking the value when the Cophenetic and Silhouette scores began to drop. Our analysis was guided by (https://htmlpreview.github.io/?https://github.com/sqjin/CellChat/blob/master/tutorial/CellChat-vignette.html) and further details can be found in our source code.

### Bone marrow decalcification

Femoral head samples were fixed for 24 hours in 4% paraformaldehyde (Electron Microscopy Sciences, 15710-S) in PBS at 4°C. Samples were washed 3 times for 30 minutes in PBS at 4°C. Samples were decalcified in 0.5M EDTA at pH8 (Invitrogen, 15575-038) for 8 days. The EDTA was changed every 2 days. After decalcification, samples were washed 5 times for 1 hour in PBS and fixed for 24 hours in 4% paraformaldehyde at 4°C. Samples were processed and embedded in paraffin by the Children’s Hospital of Philadelphia’s pathology core. The AML and NSM samples were obtained from the UPenn Department of Pathology and were previously decalcified using the following protocol: Bone marrow biopsies were fixed in buffered zinc formalin for a minimum of 1 hour. Bone marrow biopsies were then decalcified in Formical-4 (StatLab, SKU #: 1214-1) for 2 hours. Samples were rinsed with running water for a minimum of 10 minutes. Samples were placed in neutral buffered formalin and submitted for paraffin embedding.

### CODEX antibody conjugation

Akoya antibodies were purchased pre-conjugated to their respective CODEX Barcode (Supplemental Table S5). All other antibodies were custom conjugated to their respective CODEX barcode (Supplemental Table S5) according to Akoya’s PhenoCycler-Fusion user guide using the antibody conjugation kit (Akoya, 7000009) following manufacturer’s protocol. Briefly, 50 μg of carrier-free antibodies (Supplemental Table S5) were concentrated by centrifugation in 50kDa MWCO filters (EMD Millipore, UFC505096) and incubated in the antibody disulfide reduction master mix for 30 minutes. After buffer exchange of the antibodies to conjugation solution by centrifugation, addition of conjugation solution and centrifugation, respective CODEX barcodes resuspended in conjugation solution were added to the concentrated antibody and incubated for 2 hours at room temperature. Conjugated antibodies were purified by 3 buffer exchanges with purification solution. 100 μl of antibody storage buffer was added to the concentrated purified antibodies.

### CODEX staining

CODEX staining was done using the sample kit for PhenoCycler-Fusion (Akoya, 7000017) according to Akoya’s PhenoCycler-Fusion user guide with modifications to include a photobleaching step and overnight incubation in antibodies at 4°C. FFPE samples were sectioned at 5 μm thickness and mounted onto charged slides (Leica, 3800080). Sample slides were baked for 3 hours at 65°C. Sample slides were deparaffinized in Histochoice clearing agent (VWR, H103-4L) and rehydrated in a graded series of ethanol (2 times 100%, 90%, 70%, 50%, 30% and 2 times ddH2O). Antigen retrieval was performed in 1x citrate buffer (Sigma, C9999) with a pressure cooker for 20 minutes. After equilibrating to room temperature, sample slides were washed 2 times with ddH2O and submerged in a petri dish containing 4.5% H2O2 and 20mM NaOH in PBS (bleaching solution) for photobleaching. The petri dish was sandwiched between two broad-spectrum LED light sources for 45 minutes at 4°C. After 45 minutes, sample slides were transferred to a new petri dish with fresh bleaching solution and photobleached for another 45 minutes at 4°C. Sample slides were washed 3 times in PBS and then 2 times in hydration buffer. Sample slides were equilibrated in staining buffer for 30 minutes and incubated in the antibodies (Supplemental Table S5) diluted in staining buffer plus N Blocker, G Blocker, J Blocker, and S Blocker overnight at 4°C. After antibody incubation, sample slides were washed 2 times in Staining Buffer and fixed for 10 minutes in 1.6% paraformaldehyde (Electron Microscopy Sciences, 15710) in storage buffer. Sample coverslips were washed 3 times in PBS and incubated in ice cold methanol for 5 minutes. After incubation in methanol, sample slides were washed 3 times in PBS and incubated in final fixative solution for 20 minutes. The sample slides were then washed 3 times in PBS and stored in storage buffer for up to one week prior to imaging.

### CODEX imaging

CODEX Reporters were prepared according to Akoya’s PhenoCycler-Fusion user guide and added to a 96-well plate. The PhenoCycler-Fusion experimental template was set up for a CODEX Run using Akoya’s PhenoCycler Experiment Disigner software according to Akoya’s PhenoCycler-Fusion user guide. Details on the order of fluorescent CODEX Barcodes and microscope exposure times, can be found on (Supplemental Table S6). The PhenoCycler-Fusion experimental run was performed using Akoya’s Fusion 1.0.5 software according to Akoya’s PhenoImager Fusion User Guide. Images were taken and pre-processed (stitching, registration, background subtraction) with Akoya’s PhenoImager Fusion microscope using default settings.

### H&E staining and registration

Slides were removed from the microscope after CODEX imaging with the PhenoCycler platform and kept in Histochoice for 3-5 days. The flow cell was gently removed and then the tissue slide was sent to CHOP’s pathology core for standard H&E staining. Imaging was performed on the PhenoCycler Fusion microscope using the brightfield mode. 40x H&Es were registered to the 20x CODEX imaging space using wsireg (https://github.com/NHPatterson/wsireg) with affine and rigid transformations, the default parameters, with the DAPI and NaK-ATPase channels as input from the CODEX image to label the nucleus and membrane (to match with hematoxylin and eosin) respectively.

### CODEX data processing

Whole-cell segmentation was performed using Mesmer^45^ for each image. To generate the necessary input of a two-channel TIFF, we used DAPI for the nuclear channel and a composite channel of CD45, Vimentin, and NaK-ATPase for the membrane channel (see source code for further details). Mean pixel intensity was extracted from each cell segmentation mask, yielding a cell x protein matrix which was carried forward for analysis in Seurat v4. Cells with very low or high raw DAPI expression (<10 or >250 on a UINT8 scale) were removed. Images were cropped if there were instances of tissue detachment during the run, or if areas had noticeable artifactual staining. All such cropping is carefully documented in the source code. Finally, we computed the sum of all the marker expression in each cell and removed cells falling into the 99^th^ percentile of total expression to reduce the impact of reporter precipitates and similar artifacts. One Seurat object was created for each sample using this method. Each sample was then internally normalized using Seurat’s implementation of centered log ratio (CLR) which is recommended for analysis of protein expression as in CITE-Seq data. Samples were then combined into one object and integrated using reciprocal PCA (RPCA) in Seurat, this combined object was then scaled, and PCA, UMAP, and clustering was performed.

### CODEX cell type annotation

Clusters in the integrated Seurat object containing all 12 NBM samples were annotated based on the marker expression of each cluster. Clusters that could not be easily annotated were subjected to a second round of subclustering of just those ambiguous clusters and were resolved. To address the issue of lateral spillover, the concept that bright expression from one cell type may ‘spill’ into its neighbor thereby causing that cell to be mislabeled, we subjected each cluster to an additional round of subclustering, after which we visualized the subclusters on the image itself using cell phenotype maps in Napari – clusters which represented lateral spillover artifacts were corrected to the appropriate cell type. Lastly, to ensure maximum annotation accuracy, any cell labels which had been changed through this correction process were subjected to one final round of subclustering, where spillover artifacts, if present, were corrected. In some cases, we did not observe a clear cluster for a certain cell type, and we used manual gating to distinguish them, such as immunophenotypic HSCs, CLPs, and GMPs. We found highly autofluorescent cells which we used mast cell tryptase (MCT) to visualize because of the relative scarcity of bone marrow mast cells of ∼0.01%^69^, and therefore we considered cells positive for this marker to be autofluorescent and were removed from analysis. We detail the markers and strategy used for cell type annotation in Supplemental Table S4, and the source code documents the clustering/subclustering process in a step-by-step manner. Cells were assigned a coarse(level 1) and detailed (level 2) type annotation. This process is further detailed in the source code.

### CODEX neighborhood analysis

Neighborhood analysis was performed as previously described^46^ using the level 2 cell type annotations for NBM, AML, and NSM samples. AML samples had the additional cell type label of NPM1 mutant blasts which of course was not represented in either NBM or NSM samples. The ten nearest neighbors of each cell were grouped into windows, which were clustered using k-means with k=15 to get neighborhood assignments. Cell type enrichment was calculated as previously described^46^.

### NPM1 mutant cell classification and bone distance calculation

NPM1 mutant cells were identified using machine learning (Random Trees (RT) classifier) in QuPath^70^. Mutant cells were defined as those which expressed NPM1C but not CD141, as CD141 mature myeloid cells stained with the NPM1C mutant-specific antibody even in negative staging marrow controls. WT cell annotation in diagnostic AML samples was aided with visualization of lymphoid markers such as CD3E or mesenchymal markers such as FOXC1. Approximately 100 manually identified mutant and wild-type cells each were annotated using the brush tool. The cell segmentation mask defined by Mesmer was imported using custom scripts. Using QuPath, for each marker we calculated mean, standard deviation, min, max, Haralick Features (min-NaN, max-NaN, distance-1, bins-32), and the default QuPath morphologic features for each cell detection object. We then trained and applied an RT classifier to all cells from 5 samples and assigned a binary classification of NPM1_Mutant or NPM1_WT to each cell. Classification was manually evaluated in collaboration with clinical hematopathologists, and additional informative annotations were added in an iterative, human-in-the-loop fashion until the classification achieved optimal performance. Detection measurements were exported and merged with the default Seurat analysis by matching cells to their nearest spatial coordinates. Spatial analysis with respect to bone was performed in the *spatstat* package using the same approach as the healthy neighborhood distance to bone analysis.

### Negative staging marrow and AML cell type classification

AML and NSM samples were computationally processed the same as NBM samples until the cell type annotation step. Instead of using clustering and manual annotation, we used automated label transfer using RPCA in Seurat v4 with default parameters to map the NSM and AML datasets onto our NBM atlas and predict their cell labels. For comparison of NSM and AML myeloid classification proportions, we considered cells which mapped to GMP, GMP/Myeloblast, Early Myeloid Progenitor, Intermediate Myeloid, Monocyte, Non-Classical Monocyte, pDC, or Macrophages.

### Mesenchymal cell fluorescence assisted cell sorting

Femoral head samples from THA patients were digested as described above and subjected to RBC depletion and CD45 depletion using MACS (STEMCELL cat #18170 and #17898). Antibodies for FACS were purchased from BioLegend unless otherwise specified. Anti-LEPR was purchased from R&D systems (Clone # 52263, catalog: FAB867P). Isolated cells were stained for 15-30 minutes using anti-CD90-Bv650(Clone 5E10), anti-CD38-Bv785 (Clone HIT2), anti-CD45-FITC(Clone HI30), anti-LEPR-PE (Clone # 52263,R&D Systems), anti-Podoplanin-PE/Dazzle (Clone NC-08), anti-CD56-PE/Cy7 (Clone 5.1H11), anti-VE-Cadherin-Alexa Fluor 647 (Clone BV9), and anti-CD235ab-APC/Fire (Clone HIR2). Cells were spun down and resuspended in PBS + 2% FBS containing DAPI to exclude dead cells. Fluorescence minus one controls (which included all antibodies except the one of interest) were used to confirm the gating strategy, and all mesenchymal cells were gated as CD45-DAPI-CD38lo CD235ab-, hereby referred to as hem-negative. Fibro-MSCs were defined as hem-negative PDPN+ VE-Cadherin-, Osteolineage cells were defined as hem-negative PDPN-VE-Cadherin-CD56+, THY1+ MSC were defined as hem-negative PDPN-VE-Cadherin-CD56-LEPR+ CD90+ and Adipo-MSC were defined as PDPN-VE-Cadherin-CD56-LEPR+ CD90-. Cells were directly sorted into MesenCult Proliferation Kit (STEMCELL, Catalog #05411) growth media containing 25mM HEPES and cultured or used directly for downstream experiments.

### Proliferation/Cell culturing

Sorted cells were seeded in MesenCult Proliferation medium (StemCell, Catalog # 05411) for passaging. The population doubling was calculated as 3.32*log((cell count at passage)/(cells seeded)). The cumulative population doubling is reported. The cumulative population doubling for the four cell types over eight passages was analyzed by a 2-way ANOVA followed by a Bonferroni post hoc test comparing each cell type at each passage to Fibro-MSCs.

### CFU-F Assays

Cells were seeded immediately after fluorescence assisted cell sorting at 145 cells/cm^2^ in a 6-well or 12-well cell culture treated dish and maintained in MesenCult Proliferation Kit (STEMCELL, Human; Catalog #05411), until colonies could be visualized under the microscope (8-12 days). Cells were washed in PBS and stained with crystal violet (3%). Colonies were identified and manually counted as clusters with greater than 30 cells forming a tight grouping.

### Differentiation potential assessment

Sorted Fibro-MSCs were expanded in culture before plating for differentiation assays to achieve the necessary input cell number per patient. At 90% confluence, growth medium was switched to osteogenic induction medium (AlphaMEM with 10 % FBS, 1% P/S, 0.1 mM LAA, 1 mM L-glutamine, 10^-8^ M DMSO, and 1.8 mM KH_2_PO_4_) or adipogenic induction medium (AlphaMEM with 10 % FBS, 1% P/S, 0.1 mM LAA, 1 mM L-glutamine, 0.5 mM isobutylmethylxanthin, 60 µM indomethacin, 0.5 µM hydrocortisone, and 10 µg/mL insulin). Cells were harvested at day 14 for qRT-PCR analysis and at day 28 for Alizarin Red (osteogenic), Oil Red O (adipogenic), or Alcian Blue staining. For qPCR analysis, total RNA was isolated using a phenol-chloroform extraction. One μg RNA was used for reverse transcription (High-Capacity cDNA Reverse Transcription Kit, Applied Biosystem). qPCR was performed using Power SYBR™ Green PCR Master Mix. The following primers were used:

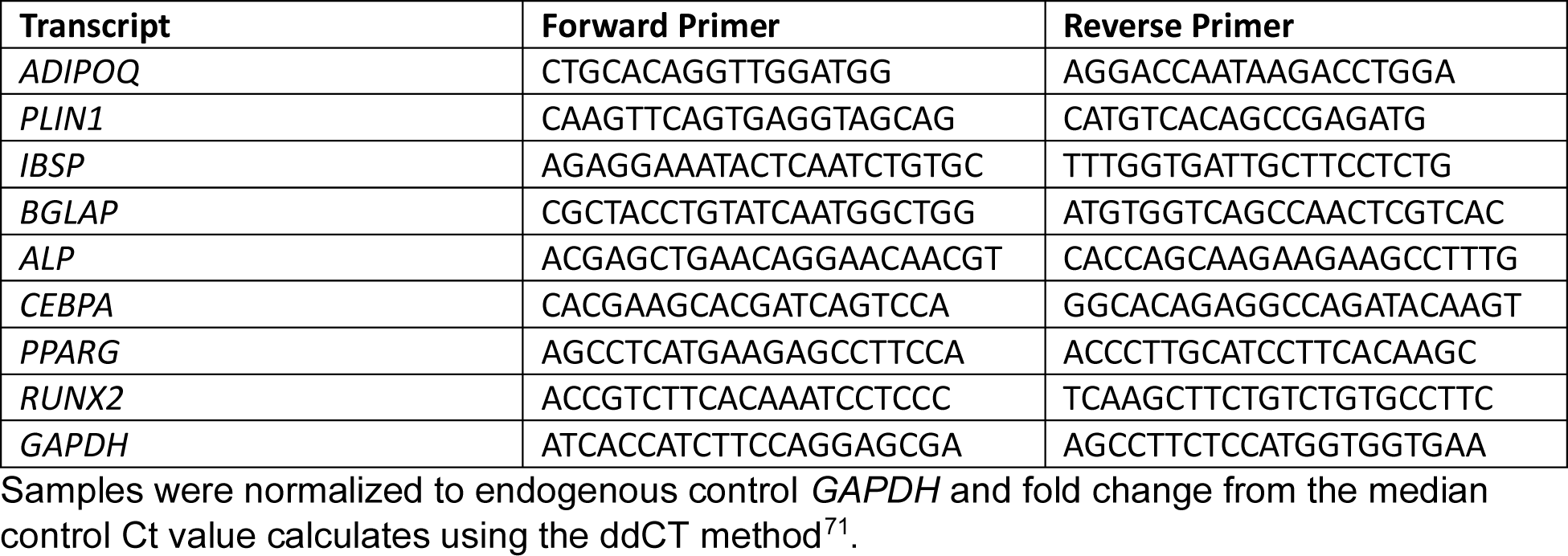

### Annotation of microenvironmental structures

Structures for proximity analysis were labeled by manual annotation and thresholding. Bone was labeled manually based on morphology and a combination of CD56 expression, exclusion of vessels, and correlation with the corresponding H&E image. Adipocytes were labeled using CD146 and morphology, with VE-Cadherin-positive arterioles and sinusoids excluded. Structures with characteristic ASMA+ cells surrounding a layer of VECAD+ CXCL12+ endothelial cells were labeled as arterioles, while vessels with VECAD/CD34 expression but not ASMA or CXCL12 were labeled as sinusoids. CXCL12+ stroma and CD163+ macrophages were labeled using manual thresholding on a per-image basis. Macrophages were considered a structure due to their considerable cytoplasmic projections which were not well captured by traditional nuclear segmentation-based approaches. Stroma, macrophages, arterioles, and bone were labeled across each entire image. Adipocytes and sinusoids were prohibitively abundant to label across the entire image, and thus large ROIs with hundreds of each structure were labeled per each image.

### Structural proximity analysis

We measured the proximity to microenvironmental structures (adipocyte, arteriolar, sinusoid, bone, macrophage, stroma) from different cell types, neighborhoods and structures using the median distance and applying a Poisson point process model. The key properties of the Poisson point process include the independence of events within non-overlapping regions and the probability of occurrence of events following a Poisson distribution. This makes the Poisson point process particularly useful for modeling rare cells.

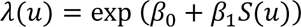

The model is applied to our data where *u* is x-y coordinate of cell, λ(*u*) is probability of cell, β_0_ is the population of cells to optimize, β_1_ is decay rate of *S*(*u*) to optimize, *S*(*u*) is distance to the structure. The value of λ(*u*) is 0 when there are no cells at position *u*, and 1 if there is a cell. When fitting the data, λ(*u*) ranges from 0 to 1. β_0_ is highly correlated to the population size, which can be counted directly. And β_1_ is what we are interested in, it represents the proximity. A small β_1_value indicates that the distance is close. The Poisson point process model is implemented in the spatstat R packages. We used the “ppm” function from that package to calculate the β_0_ and β_1_. We quantified spatial proximity using annotation mask images and a cells table that includes cell locations and labels (cell types and neighborhoods). To extract contour points for the structures, we used the cv2 package in Python. We then randomly sampled 10% of these points to create a spatial point pattern for the structures. To prevent overemphasis of outliers, we excluded rare cell types from our analysis. Specifically, if a cell type has a population of less than 10 in a given sample, it is excluded from that sample’s rank calculation. Additionally, if there are fewer than 6 samples for a particular cell type after all calculations have been completed, that cell type is also excluded from our analysis. We utilized rank proximity as our final metric, rather than physical distance, because the sizes of different marrow cavities may differ and an internally normalized metric was more appropriate. To determine the proximity of cell types, neighborhoods or structures to structures, we used a normalized rank calculation. This involved ranking the - β_1_ value for cells in comparison in every sample and scaling the ranking number to a range of 0-1 within each sample. We conducted a one-sided permutation test without replacement to determine if the cell types, neighborhoods or structures are significantly close to the structure. We permuted the labels for 100 times and calculated their median distance across the 100 permutations. The proportion of distances smaller than that of the cell type is the p-value. We used Stouffer’s method for calculation of p-values for cell types aggregated from the permutation testing p-values.

### Figure making and plotting

All figures were generated using R v4.0.5 and subsequently edited in Adobe Illustrator. Napari (doi:10.5281/zenodo.3555620) was used for generation of CODEX image figures, including fluorescent CODEX images, masks, and H&E images. Images in the manuscript figures are contrast-adjusted for illustrative purposes. For images, in certain cases Adobe Photoshop was used to globally adjust brightness or contrast for visualization purposes. GraphPad Prism v9 was used to plot Supplemental Figure 2D and perform statistical testing in Figure 2E and 2F.

## Data and code availability statement

Raw sequencing data will be deposited to SRA and processed scRNA-Seq data to Gene Expression Omnibus (GEO). CODEX data will be made available through HUBMAP’s data portal and visualization applications. Source code will be made public upon publication at gitbub, and in the meantime will be made available to reviewers upon request. Furthermore, both scRNA-Seq and CODEX data are available to explore interactively through Vitessce (https://github.com/vitessce/vitessce) widgets hosted at cscb.research.chop.edu/vitessce/demo/build (CODEX data) and cscb.research.chop.edu/vitessce/demo-seq/build (scRNA-Seq data).

## Supplemental Tables

**Supplemental Table S1** – Table summarizing the known clinical information for NBM/AML/NSM samples.

**Supplemental Table S2** – List of differentially expressed genes for all annotated cell types. A Wilcoxon rank-sum test was used comparing normalized gene expression in each cell type against all other cell types, and the default 0.25 threshold of log2-fold change was applied.

**Supplemental Table S3** – Results of the CellChat analysis in Figure 3. Each significant ligand-receptor pair is included with information including the source, target, significance, and the pathway the interaction falls into.

**Supplemental Table S4** – Summary of the cell types detected using CODEX multiplexed imaging and what markers used to annotate the different cell types.

**Supplemental Table S5** – List of antibodies used in the study, the vendors, clone information, and other relevant information about antibodies to allow for replication of our panel.

**Supplemental Table S6** – Imaging parameters including the exposure times and cycle / channel assignment for each antibody in the panel.

**Supplemental Table S7** – Comprehensive metrics for the structural analysis. Coefficient 1 is β_0_ and correlated with population size, and coefficient 2 is β_1_ and corresponds to the distance to the structure. P-value is derived from permutation testing on a per-sample basis. Both rank and normalized rank (1 is not proximal, 0 is the most proximal) are included. We computed the variability in distance as the interquartile range (75^th^ minus 25^th^ percentile of distance across cells measured in microns).

## Supplemental Figures

**Supplemental Figure 1 – Characteristics of our updated scRNA-Seq human bone marrow atlas.**

**A**) A representative image of femoral head samples received as surgical waste from total hip arthroplasty surgeries prior to enzymatic digestion, showing that there is ample visible red marrow in many of the samples. The area immediately next to the areas collected for sequencing and histology analysis was subjected to microCT analysis at a 6-μm isotropic voxel size to show the normal trabecular bone structure (circle). **B)** Diagram showing femoral head enzymatic digestion protocol. **C)** Violin plots showing the final atlas distribution of unique expressed genes, sequencing depth (UMI counts, y-axis cut at 99^th^ percentile), and percentage of reads mapping to the mitochondrial genome after filtering out low quality cells. **D)** UMAP showing the inflammatory response signature score of cells in the atlas. AUCell was used to calculate the gene set enrichment score for the GSEA Hallmark Inflammatory Response term. **E)** Normalized gene expression is projected onto UMAPs to visualize expression of key marker genes highlighting hematopoietic and non-hematopoietic cell types in the data – *CXCL12* – stromal, *NCAM1* – osteolineage, *CDH5* – endothelial, *PTPRC*-hematopoietic, *MZB1* – plasma cell, *CSF3R* – granulocyte. **F)** Comparison of stromal cell content in a major published bone marrow atlas compared to our study. The annotated bone marrow Seurat object from the Azimuth database was downloaded and the existing cluster annotations were visualized in UMAP space. The circled regions represent the non-hematopoietic bone marrow fractions captured in published data compared to our updated atlas designed to capture non-hematopoietic cells.

**Supplemental Figure S2 – Quality control, selected expression profiles, and frequencies of non-hematopoietic cell subsets.**

**A)** Violin plots showing the non-hematopoietic cell distribution of unique expressed genes, sequencing depth (UMI counts), and percentage of reads mapped to the mitochondrial genome. RNAlo MSCs were removed from subsequent analysis based on these metrics. **B)** Dotplot showing gene expression of additional canonical marker genes in bone marrow non-hematopoietic cells **C)** Day 28 micrographs of adipogenic, osteogenic and chondrogenic differentiation. Adipogenic differentiation was assessed by Oil Red O staining, osteogenic by Alizarin Red staining, and chondrogenic by Alcian blue staining. **D)** Day 14 qPCR comparing RNA expression following differentiation of cultured Fibro-MSCs using adipogenic or osteogenic medium. Control samples were taken from Day 0 of the differentiation. **E)** Stacked bar plot showing the per-sample frequency of MSC subpopulations. **F)** UMAP plots computed on just endothelial cells with overlaid expression of *PDPN, LYVE1,* and *PROX1*, which are markers of lymphatic endothelial cells. **G)** Stacked bar plots showing the per-sample frequency of endothelial cell types.

**Supplemental Figure S3 – Signaling pathways comprising intercellular communication modules.**

**A)** Gating strategy used to sort MSC subpopulations in Figure 2E-F. Sorting markers were selected based on scRNA-Seq. **B)** Violin plot showing normalized expression of genes coding for markers used in cell sorting to distinguish MSC subsets. **C)** UMAP plot of the entire scRNA-Seq atlas with overlaid normalized *CSF3* expression across all cell types in the atlas. **D)** Heatmap showing the contribution of each specific signaling pathway to the inferred cellular communication modules from Figure 3. Contribution scores were calculated as previously described in Jin et al., *Nature Communications*, 2021.

**Supplemental Figure S4 – CODEX cell typing validation and cell phenotype maps.**

**A)** CODEX images with selected markers and corresponding cell phenotype maps (CPMs) showing appropriate co-labeling of certain markers and associated cell phenotype maps with registered H&E images. Grey cell masks refer to all other segmented cells in the final analysis (imaging artifacts and Mast Cell Tryptase (MCT)+ autofluorescent cells removed). MCT is used as a marker of autofluorescence in these images (See Methods). Further details on marker combinations used to define each cell type can be found in Supplemental Table S4. Registered H&E images of the exact imaged region is also provided. **B)** CODEX image of Podoplanin+ CXCL12+ Fibro-MSC corresponding to our scRNA-Seq data demonstrating that the rare cells are found in the region where the bone detached from the slide (boundary manually illustrated in grey, drawn based on H&E imagesand morphology). MCT is included as it labeled highly autofluorescent cells. **C)** CODEX image showing an Osteo-MSC and osteoblast at the bone boundary, where the bone detached from the slide (boundary manually illustrated in grey, drawn based on H&E staining and morphology). **D)** CODEX image of macrophages showing that CD163 better labels cytoplasmic projections but bone marrow macrophages label for both CD68 and CD163. MCT is included to label autofluorescent cells.

**Supplemental Figure S5 – CODEX reveals a relatively hyperoxygenated peri-arteriolar/peri-endosteal niche.**

**A)** Boxplot showing each neighborhood ranked by its proximity to manually annotated bone in each sample, and the normalized rank was computed for each comparison, such that a value of 0 means most proximal and 1 means least proximal. **B)** CODEX images shown with selected markers and paired neighborhood masks demonstrating the capturing of spatial organization of the tissue with respect to both hematopoietic and non-hematopoietic cell types. Neighborhoods with the same annotation (e.g. Erythroid 1 (CN13) and Erythroid 2 (CN15)) were combined for visualization purposes. **C)** Violin plots showing centered log ratio (CLR) HIF1a expression across all cell types.

**Supplemental Figure S6 – Structural analysis annotations and result summary.**

**A)** Paired CODEX images and overlaid masks with representative examples of the structural masks compared to the actual fluorescent images. The sinusoid and adipocyte annotations are shown in the same region of interest. **B)** Chord diagram showing only significant interactions (p<0.05, Stouffer’s method for meta-analysis of permutation test p-values across all 12 samples), chord thickness is proportional to the negative log_10_ of the median normalized rank + 0.000000000001.

**Supplemental Figure S7 – CODEX enables classification and downstream analysis of *NPM1* mutant blasts.**

**A)** Violin plot of CODEX CLR-normalized protein expression showing the non-specific expression of the mutant-specific NPM1C antibody in negative staging marrow samples. **B)** CODEX images showing that CD141, but not early myeloid or erythroid markers (MPO and GATA1) almost completely co-stains with mutant-specific NPM1C in a representative negative staging marrow sample, showing that exclusion of these CD141+ cells enables use of the antibody to identify true mutant cells. **C)** Chart showing the detected NPM1c mutant percentage in each sample compared to the clinical variant allele frequency for NPM1c. **D)** CODEX fluorescent image of a diagnostic AML sample showing MSCs and *NPM1* mutant blasts. White arrows point to FOXC1+ CD271+ MSCs. **E)** Boxplot showing the relative proximity to bone (normalized rank by AML/NSM neighborhood, where 0 is the closest in a sample and 1 is the furthest), with the leukemic neighborhoods highlighted in magenta and CN7, the neighborhood corresponding to the healthy early myeloid progenitor niche, highlighted in blue. All other neighborhoods are green. **F)** Violin plots of myeloid cell types in AML and negative staging marrow samples showing that *NPM1* mutant blasts have low HIF1a levels consistent with an early myeloid progenitor phenotype. **G-H)** BCL2 and Complex IV expression levels in *NPM1* mutant blasts. Wilcox Rank Sum test was performed in Seurat to compare between diagnostic samples and post-therapy samples.

## Acknowledgments

The authors thank the Children’s Hospital of Philadelphia Pathology Core for their considerable technical expertise and assistance in handling human tissue sections and the Research Information Services for providing computing support. We thank the many clinical contributors who assisted in procuring primary human samples through Penn Presbyterian Medical Center. Lastly, we also thank the University of Pennsylvania Cytomics and Cell Sorting Shared Resource Laboratory for their assistance in sorting primary MSC samples.

This work was supported by National Institutes of Health of United States of America grants U54 HL156090 and U54HL165442 (to KT), R01AG069401 (to LQ), and P30AR069619 (to The University of Pennsylvania). MD was supported by NIH NIDDK T32DK007314. ShB was supported by NIH T32 GM007170, T32 HL007439, and F30CA277965.

## Relative Contributions

KT, LQ, and ShB conceptualized and designed the project. ShB, KJA, MD, AT, ChiC, and YL performed the data generation and experiments. ShB, KJA, MD, JS, GD, JH, IZ, and TI performed analysis and wrote software and made figures. ShB, KJA, MD, JX, ChaC, VP, IM, MC, and CN, LQ, and KT provided critical conceptual contributions and technical expertise. CN, SiB, and and OS assisted with reagent validation and provided clinical samples. ShB, KJA, MD, LQ, and KT wrote the manuscript.

